# Discovery of highly potent pancoronavirus fusion inhibitors that also effectively inhibit COVID-19 variants from the UK (Alpha), South Africa (Beta), and India (Delta)

**DOI:** 10.1101/2021.09.03.458877

**Authors:** Francesca Curreli, Shahad Ahmed, Sofia M. B. Victor, Aleksandra Drelich, Siva S. Panda, Andrea Altieri, Alexander V. Kurkin, Chien-Te K. Tseng, Christopher D. Hillyer, Asim K. Debnath

**Author notes:** Corresponding authors Francesca Curreli- Laboratory of Molecular Modeling & Drug Design, Lindsley F. Kimball Research Institute, New York Blood Center, New York, New York, USA, Asim K Debnath- Laboratory of Molecular Modeling & Drug Design, Lindsley F. Kimball Research Institute, New York Blood Center, New York, New York, USA.

## Abstract

We report here the discovery of several highly potent small molecules that showed low nM potency against SARS-CoV (IC_50_: as low as 13 nM), SARS-CoV-2 (IC_50_: as low as 23 nM), and MERS-CoV (IC_50_: as low as 76 nM) in pseudovirus based assays with excellent selectivity indices (SI: as high as > 5000) demonstrating their pancoronavirus inhibition. Some compounds also show 100% inhibition of CPE (IC_100_) at 1.25 µM against an authentic SARS-CoV-2 (US_WA-1/2020). Furthermore, the most active inhibitors also potently inhibited variants of concerns (VOCs), such as the UK (B.1.1.7), South Africa (B.1.351), and Delta variant (B.1.617.2), originated in India. We confirmed that one of the potent inhibitors binds to the prefusion spike protein trimer of SARS-CoV-2 by SPR. Besides, we showed that they inhibit virus-mediated cell-cell fusion. The ADME data of one of the most active inhibitors, NBCoV1, show drug-like properties. In vivo PK of NBCoV1 in rats demonstrated excellent half-life (t1/2) of 11.3 h, mean resident time (MRT) of 14.2 h, and oral bioavailability. We expect the lead inhibitors to pave the way for further development to preclinical and clinical candidates.

## INTRODUCTION

The outbreak of coronavirus disease 2019 (COVID-19) caused by the novel coronavirus SARS-CoV-2 and first reported in December 2019 in Wuhan^1^, China, has led to massive human suffering, death, and economic devastation throughout the world. Thanks to the exceptional ingenuity of academics and pharmaceutical companies, several highly effective vaccines against SARS-CoV-2, the virus that causes COVID-19, were developed in record time, including one vaccine (Pfizer-BioNTech) approved recently and two vaccines (Moderna and Janssen) were approved under emergency authorization by the U.S. Food and Drug Administration (FDA). Despite this breakthrough development, effective vaccines may not reach all individuals worldwide. In addition, vaccine hesitancy is proving to be a major roadblock to getting populations fully vaccinated in many countries^2–4^. According to the U.S. Census Bureau’s Household Pulse Survey, up to 32% of the U.S. population is reluctant to get vaccinated against SARS-CoV-2. Furthermore,“breakthrough” SARS-CoV-2 infections have been reported among people who were fully vaccinated^5, 6^. Therefore, the development of novel drugs to treat or prevent infection is urgently needed. Coronaviruses (CoVs) are positive-sense, enveloped, single-stranded RNA viruses in the family Coronaviridae, which includes two other CoVs that caused major outbreaks in recent years: Severe Acute Respiratory Syndrome coronavirus (SARS-CoV; originated in China in 2003) and the Middle East Respiratory Syndrome coronavirus (MERS-CoV; originated in Saudi Arabia in 2012). Although these outbreaks were severe, they have been eclipsed by the current COVID-19 pandemic. According to data from the Johns Hopkins Coronavirus Resource Center, as of September 10, 2021, there were more than 223 million cases of COVID-19 and 4.6 million deaths due to COVID-19 globally. In the U.S., over 42 million cases and more than 655,000 deaths have been reported.

Although one broad-spectrum antiviral, remdesivir, was approved by the FDA to be repurposed for emergency use against COVID-19, it has shown limited efficacy in most clinical settings. Several monoclonal antibody-based therapies, including those from Regeneron Pharmaceuticals and Eli Lilly, were also approved by the FDA for use against COVID-19, but their high cost and limited accessibility are prohibitive to a significant part of the world. Furthermore, most of the approved vaccines and antibody-based therapies show substantial loss of potency against SARS-CoV-2 variants that emerged in the U.K. (Alpha variant; B.1.1.7), South Africa (Beta variant; B.1.351, and Brazil (Gamma variant; P.1) are now spreading across the globe^7–10^. Recently, the Delta variant (B.1.617.2)^6, 11, 12^, identified initially in India, is spreading like wildfire. While vaccines are critical in preventing infection and severe illness, therapeutic drugs play a crucial role in combatting the disease in individuals who get infected. Therapeutic drugs may shorten symptomatic disease or prevent serious illness, hospitalization, or death in patients with SARS-CoV-2 infection. There is currently no FDA-approved drug that targets the host-cell entry/fusion of CoVs. Therefore, the development of highly potent novel drugs with pancoronavirus activity with minimal toxicity is urgently needed. To emphasize the value of therapeutics, it is noteworthy that 245 antivirals are currently under development against COVID-19 (https://www.bio.org/policy/human-health/vaccines-biodefense/coronavirus/pipeline-tracker ).

Like all enveloped viruses, CoVs begin their life cycle with entry into a host cell, which is initiated by the trimeric spike surface protein (S)^13, 14^. The S protein is cleaved into S1 and S2 subunits by furin-like proteases once it binds to a host cell. The S1 subunit of the S protein attaches to a host-cell receptor through its receptor-binding domain (RBD). The receptor for SARS-CoV and SARS-CoV-2 is angiotensin-converting enzyme 2 (ACE2)^15, 16^, whereas the receptor for MERS-CoV is dipeptidyl peptidase 4 (DPP4; also termed CD26)^17^. After the S1 subunit binds a receptor, the fusion protein (FP) of the S2 subunit inserts into the cell membrane and triggers the heptad repeat region 1 (HR1) domain of S2 to form a coiled-coil trimer. The heptad repeat region 2 (HR2) domain of S2 then binds to a hydrophobic groove in the HR1 trimer in an antiparallel manner, creating a six-helix bundle (6-HB) structure. This process brings the viral membrane and the host-cell membrane together for virus–cell fusion^18^, a critical step for virus entry into host cells. The mechanism of fusion for the S protein of CoVs, including MERS-CoV, is like that for other class I membrane fusion proteins in viruses such as influenza virus, human immunodeficiency virus (HIV), and Ebola virus^13, 19^. Some distinctions exist, however, including the larger size, double cleavage site, and long 6-HB structure of the CoV S protein^13^.

Because of its exposure on the surface of the mature virus particle, the S protein is the primary target for neutralizing antibodies and vaccines. Both the S1 subunit, primarily the RBD domain, and the S2 subunit, especially the HR1 domain, have been targeted for novel drug design^20–28^. The RBD of the S1 domain is not well conserved among CoVs^29^, however. Therefore, antibodies that neutralize SARS-CoV show poor cross-reactivity with SARS-CoV-2^30, 31^. Furthermore, several mutations in the RBD domain of SARS-CoV-2 have been reported^32, 33^. Some of these mutations reduce the efficacy of antibodies, and hence the currently available vaccines^34, 35^. Therefore, the RBD may not be an ideal target for the development of novel pan-coronavirus inhibitors.

On the other hand, the membrane fusion domains located in the S2 subunit are mostly conserved and have the potential to be an ideal target for novel small-molecule and peptide-based pan-coronavirus inhibitors. Drugs that target the most conserved sites among CoVs will provide better broad-spectrum (pancoronavirus) antiviral activity^29^. These drugs will be critically important in dealing with new pandemics that are certain to emerge in the future.

Herein, we report the identification and characterization of a series of inhibitors that show highly potent pancoronavirus activity against SARS-CoV, SARS-CoV-2, and MERS-CoV. Moreover, the most active inhibitors also potently inhibit laboratory synthesized mutants, mimicking the currently circulating variants B.1.1.7 UK (Alpha), B.1.351 RSA (Beta), and B.1.617.2 India (Delta). The lead inhibitors are expected to pave the way for further development to preclinical and clinical candidates.

## RESULTS AND DISCUSSION

### Rationale of screening HIV-1 fusion inhibitors as possible pancoronavirus inhibitors

Spike protein of coronaviruses plays a critical role in virus binding to a cellular receptor, subsequent fusion, and entry into host cells, allowing the release of the genetic material for continuing its life cycle. The fusion protein (FP) inserts into the host cell membrane and triggers the formation of a coiled-coil trimer by the heptad repeat region 1 (HR1). The HR2 binds to the trimer’s hydrophobic groove in an antiparallel manner, creating a 6-HB, like what was reported for HIV-1 gp41-mediated fusion^36, 37^. However, there is no sequence homology between the HR1 and HR2 of coronaviruses spike protein with HR1 and HR2, respectively, of HIV-1 gp41. The formation of the 6-HB facilitates the virus membrane and host cell membrane to come closer for the fusion process to complete.

The mechanistic similarity between the fusion processes of SARS-CoV and HIV-1 led Jiang et al. to design peptide-based inhibitors with pan-coronavirus activity based on the HR2 domains of SARS-CoV^38^ and SARS-CoV-2 ^25^. Jiang et al. crystallized the postfusion hairpin structure of SARS-CoV-2 (6LXT)^25^ and compared that structure with the corresponding SARS-CoV structure (1WYY)^39^ reported in 2005. Remarkably, the postfusion hairpin structures were not only structurally similar but also shared critical salt bridges between the HR1 and HR2 regions^25^. Most noteworthy was the finding that in SARS-CoV-2, K947 of the HR1 domain forms a salt bridge with E1182 of the HR2 domain, while in SARS-CoV, K929 of the HR1 domain forms a salt bridge with E1163 of the HR2 domain. Furthermore, even the MERS-CoV postfusion spike structure showed a salt bridge in a similar position- K1021 forms a salt bridge with E1265^40^. We previously reported a similar salt bridge interaction in the HIV-1 gp41 hairpin structure, where K547 of the N-terminal heptad repeat region (also known as HR1) interacts with D632 of the C-terminal heptad repeat region (also known as HR2)^41^. We also reported the design of a series of highly potent, benzoic acid-based inhibitors of HIV-1 gp41 fusion, which contain a COOH group^42^.

Also, we showed through computer-based docking that the COOH group of these inhibitors might interrupt the hairpin structure formation by interacting with the K547 and snuggly fitting to the hydrophobic grove created by three NHR regions^42^. These remarkable similarities in the mechanism of virus fusion and involvement of salt bridges in forming the 6-HB in HIV-1 gp41 and coronaviruses lead us to hypothesize that this class of inhibitors may also interrupt the salt bridge formation by fitting in a cavity in the prefusion trimer of coronaviruses thus preventing the 6-HB formation and eventually the fusion of viruses to host cells. Therefore, we screened a set of nine such compounds from our stock and four new analogs (NBCoV15, NBCoV17, NBCoV28, and NBCoV34) against SARS-CoV-2, SARS-CoV, and MERS-CoV spike pseudotyped antiviral assay (**Figure 1**). We added the new four analogs to derive a comprehensive structure-activity relationship (SAR). It is noteworthy that recently peptide-based pancoronavirus fusion inhibitors were shown to possess potent inhibitory activity against HIV-1, HIV-2, and SIV^43^.

**Figure 1.**
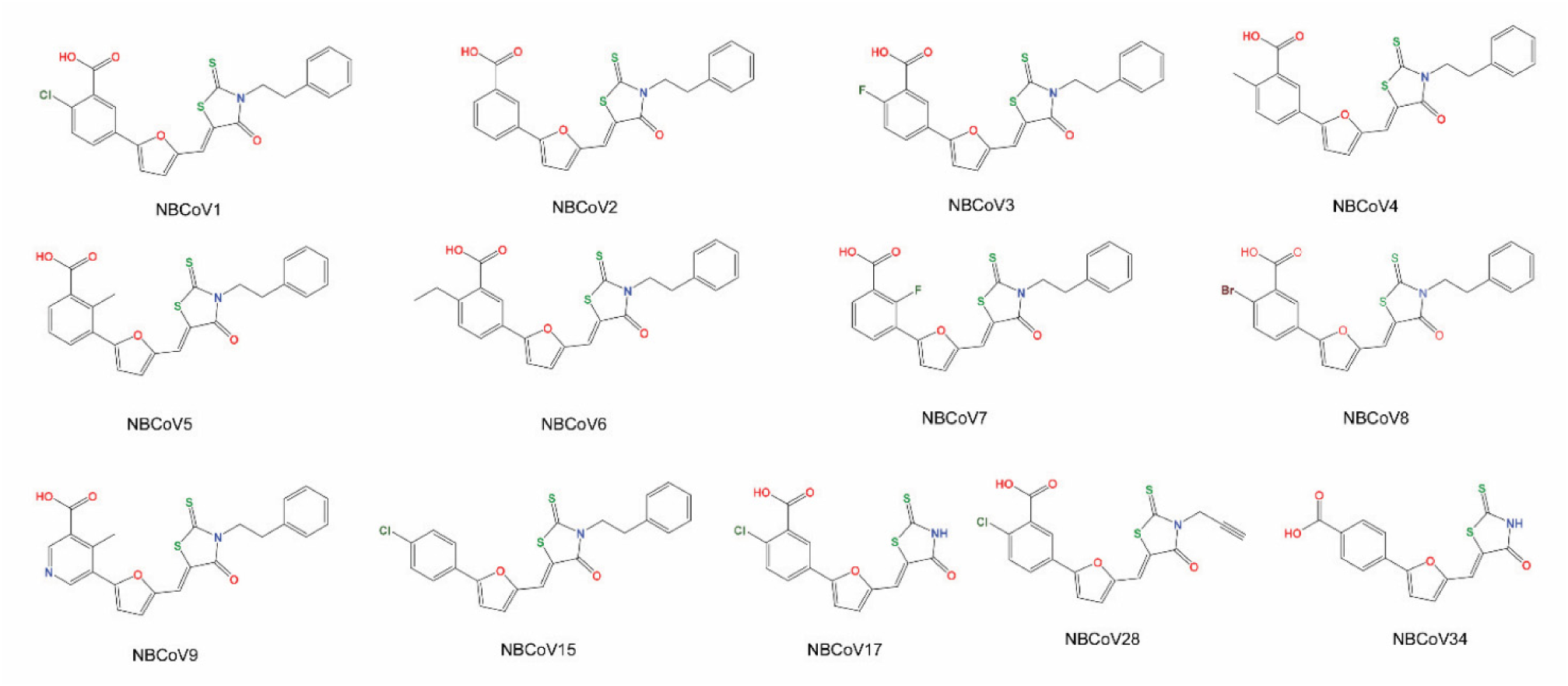
The structures of all the 5-((5-(4-chlorophenyl)furan-2-yl)methylene)-3-phenethyl-2-thioxothiazolidin-4- one tested against SARS-CoV-2, SARS-CoV, and MERS-CoV.

We are aware that the selected inhibitors contain a well-known frequent hitter scaffold, ene-rhodanine, identified as PAINS (Pan Assay INterference compoundS)^44–46^. However, instead of discarding these molecules as most likely promiscuous, we decided to demonstrate in pain-staking details that this is a privileged scaffold in the context of pancoronavirus inhibition like reported by others^47^.

### Validation of the pseudoviruses

We prepared SARS-CoV-2, SARS-CoV, and MERS-CoV pseudoviruses capable of single-cycle infection by transfecting HEK293T/17 cells with an HIV-1 Env-deleted proviral backbone plasmid pNL4-3ΔEnv-NanoLuc and the respective spike plasmid^48^. We then validated the incorporation of the spike proteins in the respective pseudoviruses by Western blot analysis. We used the SARS Spike Protein Antibody (Novus Biologicals) to immunodetect the spike protein S2 of SARS-CoV-2 and SARS-CoV pseudoviruses (**Figure 2a**). Similarly, we used MERS-coronavirus spike protein S2 polyclonal antibody (Invitrogen) to immunodetect the spike protein S2 of the MERS-CoV pseudovirus (**Figure 2b**). For SARS-CoV-2, we found a specific band at 80 kDa, which identifies the subdomain S2, and a second band at about 190 kDa, which corresponds to the full-length S protein (S1 + S2), as previously reported ^27, 49, 50^. In SARS-CoV pseudovirus lysate, the same antibody detected a lighter band at 80 kDa representing the S2 subunit and a 190 kDa band, which corresponds to the full-length of the spike S protein (**Figure 2a**). Additionally, for MERS-CoV lysates, we identified the subdomain S2 at 75 kDa and the full-length S protein at about 185 kDa (**Figure 2b**) as previously reported^30, 51^. Thus, these analyses confirmed the correct incorporation of the spike proteins in the respective pseudoviruses.

**Figure 2.**
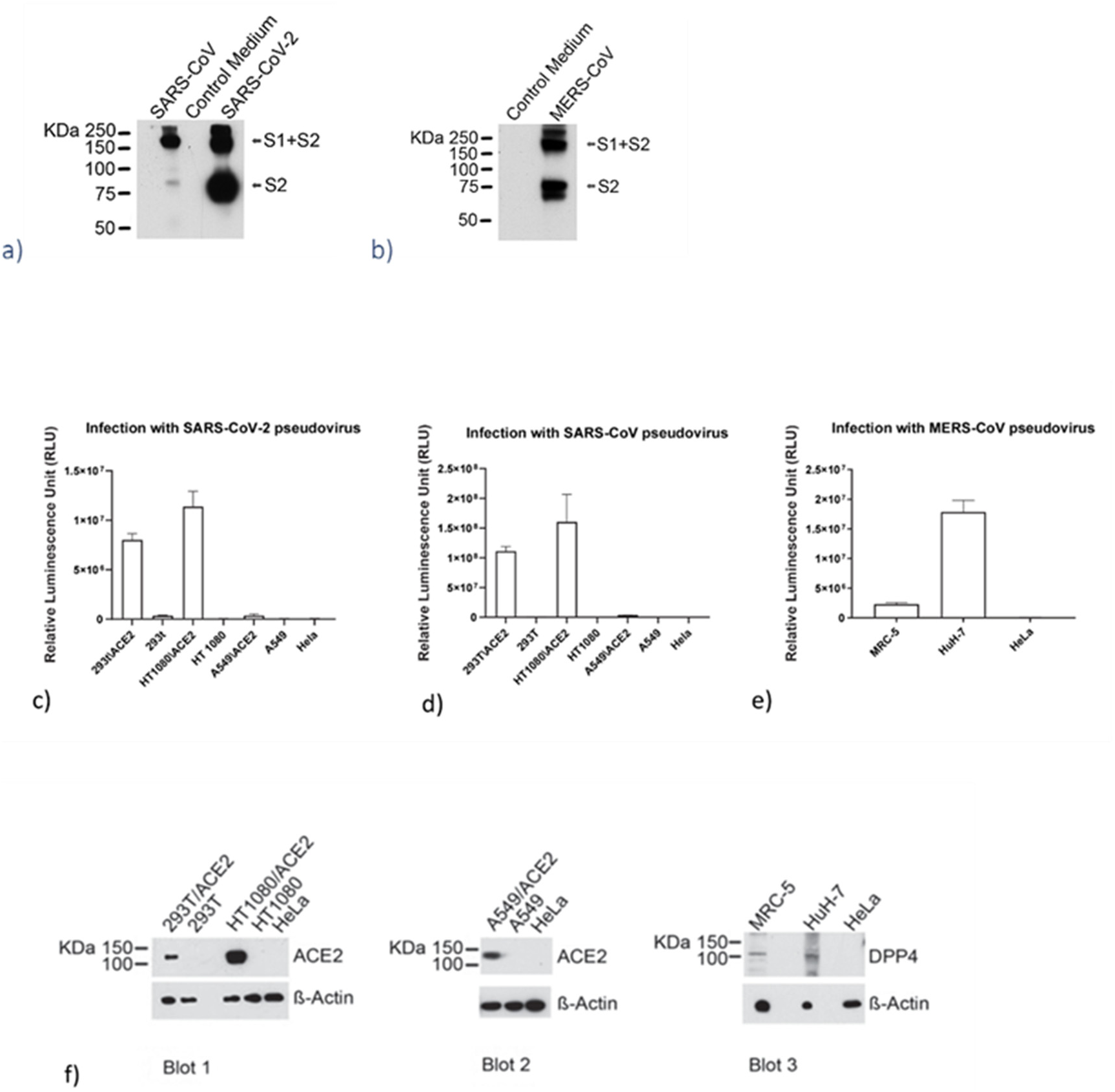
**Validation of the SARS-CoV-2, SARS-CoV, and MERS-CoV pseudoviruses and ACE2 and DPP4 expression in different cell lines.** a) lmmunoblot to validate the incorporation of the S spike protein in the SARS-CoV and SARS-CoV-2 pseudoviruses b) lmmunoblot to validate the incorporation of the S spike protein in the MERS-CoV pseudovirus Infection of cells expressing different levels of ACE2 with c) the same amounts of SARS-CoV-2 pseudovirus, d) the same amounts of SARS-CoV pseudovirus. e) Infection of cells expressing different levels of DPP4. with the same amounts of MERS-CoV pseudovirus. Columns represent the means ± standard deviations (n=4). f) lmmunoblot of cell lysates to evaluate ACE2 expression (Blot 1 and Blot 2) and DPP4 expression (Blot 3). B-Actin was used as a loading control.

Afterward, we analyzed the correlation of SARS-CoV-2 and SARS-CoV pseudoviruses infection levels with the expression of the hACE2 receptor by infecting three different cell types, which overexpress the ACE2 receptor. We used the human kidney 293T/ACE2 cells, the human fibrosarcoma HT1080/ACE2 cells, the human lung carcinoma cells A549/ACE2, and the respective parental cell types HEK293T cells, HT1080 cells, and A549 cells. We also infected HeLa cells as a control that do not express the hACE2 receptor. The cells were exposed to the same volumes of the supernatant containing the respective pseudoviruses. As expected, we found that both pseudoviruses did not infect the HeLa cells (**Figure 2c, 2d)**. Similarly, we detected a low level of infection of the parental cell lines HEK293T, HT1080, and A549 with the SARS-CoV-2 pseudovirus compared to the respective related cell types overexpressing the ACE2 receptor. The 293T/ACE2 and HT1080/ACE2 cells supported high levels of SARS-CoV-2 infection. We detected about 8×10^6^ RLU and 1.1×10^7^ RLU, respectively, which correspond to a 24-fold and 490-fold higher infection than we detected for the parental cell type HEK293T and HT1080, respectively. The infection detected for the A549/ACE2 cells was moderate (about 3.8×10^5^ RLU) compared to HT1080/ACE2 and 293T/ACE2 cells and about 13-fold higher than what we detected for the parental cell type A549. For the SARS-CoV infection study, we found similar results. In this case, the 293T/ACE2 and HT1080/ACE2 cells supported higher infection than what we detected for the parental cell types and the A549/ACE2 cell type. Then, we wanted to confirm these results by analyzing the expression levels of the ACE2 receptor in the different cell lines by western blot (**Figure 2f**). As shown in Blot 1, we found that ACE2 expression was undetectable in the parental cell lines 293T and HT1080 while it was overexpressed in HT1080/ACE2 cells. A lower amount of ACE2 was detected in the 293T/ACE2 cell lysate than the amounts detected in HT1080/ACE2 cells confirming the infection study findings (**Figure 2c,2d**). The lower infection detected in A549/ACE2 cells suggested a lower ACE2 expression in these cells. For this reason, in Blot 2 (**Figure 2f**), to visualize the ACE2 expression, we had to load a higher concentration of proteins (75 µg) and use a 2x concentration of antibodies. These data, taken together, confirmed that SARS-CoV-2 and SARS-CoV pseudoviruses infect the cells through their interaction with the ACE2 receptor.

To analyze the correlation of MERS-CoV pseudovirus infection levels with the DPP4 (CD26) receptor expression, we infected the fibroblast cell line from lung MRC-5 and hepatocyte-derived carcinoma cell line HuH-7; as a control, we also infected HeLa cells that do not express the DPP4 receptor. In this case, the cells were exposed to the same volumes of the supernatant containing the MERS-CoV pseudovirus. We found that the HuH-7 cells supported an 8.6-fold higher level of MERS-CoV infection than the MRC-5 cells. We detected about 2×10^7^ RLU and 2.3×10^6^ RLU, respectively. We observed no infection of the HeLa cells (**Figure 2e**). The expression levels of the DPP4 receptor in the two cell lines are reported in Blot 3 (**Figure 2f)**. These data, taken together, confirmed that the spike proteins were rightly incorporated in the respective pseudoviruses. It also demonstrated that these pseudoviruses infected the cells through their interaction with the respective receptors.

### Antiviral activity and cytotoxicity of the NBCoV small molecules in a pseudovirus assay

We evaluated the anti-coronavirus activity of the NBCoV small molecules by infecting the three cell types overexpressing the receptor, namely, 293T/ACE2, HT1080/ACE2, and A549/ACE2 cells, with aliquots of the SARS-CoV-2 pseudovirus, which was pretreated with escalating concentrations of the NBCoV small molecules for 30 minutes. We calculated the concentration of NBCoV small molecules required to inhibit 50% (IC_50_) of SARS-CoV-2 pseudovirus infection, and the results are reported in **Table 1.** We demonstrated that most of the NBCoV compounds inhibited SARS-CoV-2 infection with low nanomolar (nM) activity. The only exceptions were NBCoV5 which had micromolar activity (1205 ± 240 nM, 1050 ± 252 nM, and >2000 nM in 293T/ACE2, HT1080/ACE2, and A549/ACE2 cells, respectively), and NBCoV15, used as a control compound without a COOH group in the phenyl ring, which showed no antiviral activity at 2000 nM. Additionally, compounds NBCoV17, NBCoV28, and NBCoV34 at 2000 nM showed no antiviral activity as well. NBCoV1, NCoV2, and NBCoV4 were the most potent compounds. In fact, for NBCoV1, the calculated IC_50_s were in the range of 32.3-63.4 nM, and the selectivity index (SI obtained from the ratio CC_50_/IC_50_) ranged from 755 - 2755; for NBCoV2, the calculated IC_50_s were in the range of 22.8 - 58 nM, and the SIs varied from 1630-> 4000; finally, for NBCoV4 the IC_50_s were in the range of 26 -73 nM, and the SIs ranged from >1370 -> 2096. The NBCoV3 also displayed potent anti-SARS-CoV-2 activity, but the IC_50_s and SIs obtained for the three cell lines were slightly higher than those detected for NBCoV1, NCoV2, and NBCoV4 (IC_50_s: 60.1 - 120 and SIs:750 -> 1563). The remaining NBCoV compounds (NBCoV6 - NBCoV9) showed lower potency than NBCoV1 - NBCoV4, as demonstrated by the higher IC_50_s and SIs. We noticed that all the compounds displayed better activity when tested in 293T/ACE2 and HT1080/ACE2 cells than when tested in A549/ACE2 cells and consistently showed an increase of the IC_50_s and SIs. The cytotoxicity (CC_50_) of the small molecules was assessed in parallel with the inhibition assays (the data is reported in **Table 1**), and these values were used to determine the SIs. As noted, in some cases (HT1080/ACE2 and A549/ACE2 cell lines), the small molecules did not induce any apparent cell toxicity at the highest dose tested (**Table 1**). As expected, when the cells (rather than the virus) were pretreated with the NBCoV compounds for 30 minutes before infection, they did not confer any protection against SARS-CoV-2 infection even at the higher dose used in the assay (2000 nM) (**Table S1**). The data validate the hypothesis that the target of these compounds is virus-related and not cell-related.

**Table 1.** Antiviral activity of the NBCoV small molecules in the single-cycle assay in different cell lines against pseudovirus NL4-3 Env-Nanol uc/SARS-CoV-2 (IC_50),_ toxicity (CC_50),_ and selectivity index (SI).

| Compound | 293T/ACE2 cells |  |  | HT1080/ACE2 cells |  |  | A549/ACE2 cells |  |  |
| --- | --- | --- | --- | --- | --- | --- | --- | --- | --- |
|  | IC<sub>50</sub> (nM)<sup>a</sup> | CC<sub>50</sub> (μM)<sup>a</sup> | SI | IC<sub>50</sub> (nM)<sup>a</sup> | CC<sub>50</sub> (μM)<sup>a</sup> | SI | IC<sub>50</sub> (nM)<sup>a</sup> | CC<sub>50</sub> (μM)<sup>a</sup> | SI |
| NBCoV1 | 51±17 | 38.5±1 | 755 | 32.3±4.6 | 89±2 | 2755 | 63.6±4.6 | 86±8.7 | 1352 |
| NBCoV2 | 22.8±0.8 | 37.5±2 | 1630 | 25.3±0.6 | >100 | >4000 | 58±1.7 | >100 | >1724 |
| NBCoV3 | 60.1±8.5 | 45±0.5 | 750 | 64±18 | >100 | >1563 | 120±5 | >100 | >833 |
| NBCoV4 | 26±1 | 40.7±2.3 | 1565 | 47.7±16 | >100 | >2096 | 73±4.1 | >100 | >1370 |
| NBCoV5 | 1205±240 | 35±2 | 29 | 1050±252 | >100 | >95 | >2000 | >100 | N/A<sup>b</sup> |
| NBCoV6 | 185±5.8 | 40±2.6 | 216 | 245±5 | >100 | >408 | 613±72 | >100 | >163 |
| NBCoV7 | 298±60 | 45±0.4 | 151 | 290±57 | >100 | >345 | 416±25 | >100 | >240 |
| NBCoV8 | 65.8±6.2 | 38.7±1.2 | 586 | 94±9 | >100 | >1063 | 254±27 | >100 | >394 |
| NBCoV9 | 342±46 | 33.7±2.5 | 98 | 365±73 | >100 | >274 | 596±42 | >100 | >168 |
| NBCoV15 | >2000 | 64±14 | N/A<sup>b</sup> | >2000 | >100 | N/A<sup>b</sup> | N/A<sup>b</sup> | N/A<sup>b</sup> | N/A<sup>b</sup> |
| NBCoV17 | >2000 | >100 | N/A<sup>b</sup> | >2000 | >100 | N/A<sup>b</sup> | N/A<sup>b</sup> | N/A<sup>b</sup> | N/A<sup>b</sup> |
| NBCoV28 | >2000 | >100 | N/A<sup>b</sup> | >2000 | 92±3 | N/A<sup>b</sup> | N/A<sup>b</sup> | N/A<sup>b</sup> | N/A<sup>b</sup> |
| NBCoV34 | >2000 | 83±4 | N/A<sup>b</sup> | >2000 | ~100 | N/A<sup>b</sup> | N/A<sup>b</sup> | N/A<sup>b</sup> | N/A<sup>b</sup> |

<sup>a</sup>The reported IC<sub>50</sub> and CC<sub>50</sub> values represent the means ± standard deviations (n = 3).
<sup>b</sup>Not Available

To derive a structure-activity relationship (SAR), we observed that NBCoV2, which does not have any para-substituent in the carboxyphenyl ring, showed the best antiviral potency against SARS-CoV-2. The compounds (NBCoV1, NBCoV4, and NBCoV8) also showed excellent antiviral activity when there is a hydrophobic substituent at the para position of that ring. NBCoV3 has a highly electronegative fluoro atom at the para-position of the carboxyphenyl ring, and it showed somewhat reduced activity compared to NBCoV2 and NBCoV4. A bulkier hydrophobic substituent (-CH_2_CH_3_) in NBCoV6 showed slightly lower antiviral activity. Furthermore, a substituent at the ortho position in the carboxy phenyl ring did not tolerate well and showed poor antiviral activity. The introduction of nitrogen in the phenyl ring (pyridine) did not improve the antiviral activity, although the solubility of this molecule may have improved. Since we hypothesized that the COOH group of these compounds might be interacting with one of the key positively charged residues in the HR1 region to interfere with the 6-HB formation, we tested an analog devoid of the COOH group (NBCoV15). As expected, this molecule showed no antiviral potency at the highest dose tested. We extended the SAR by substituting the phenylethyl group with either no substitution in the rhodanine moiety (NBCoV17 and NBCoV34) or substituting with a smaller hydrophobic group (prop-1-yne, NBCoV28). Furthermore, the position of COOH is also critical. When the COOH group is at the para position of the phenyl ring (NBCoV34), it loses the inhibitory activity. The data, although with few compounds, clearly generated an insightful SAR.

Although NBCoV17 and NBCoV28 contain COOH and Cl groups at the same positions as NBCoV1, they did not show any activity at the highest dose tested. This data underscores the fact that a combination of electronic and appropriate hydrophobic interactions is essential for the antiviral potency of this series of molecules. The mere presence of the ene-rhodanine moiety shows no role in the inhibition process. Similarly, NBCoV34 had a carboxylic group at the para position, had no hydrophobic group attached to the NH of the rhodanine moiety, and showed no inhibition. Suppose the ene-rhodanine scaffold has any role in the antiviral potency through its promiscuous nature, as claimed. In that case, it should have shown potent antiviral activity because the rhodanine scaffold has no steric hindrance to bind non-specifically to random protein targets. Based on the above observations, we only selected the most active inhibitors to test against other coronaviruses.

To assess whether the NBCoV small molecules have pancoronavirus antiviral activity, we tested them against the SARS-CoV pseudovirus (**Table 2**). We also found that, in this case, the most potent compounds were NBCoV1 - NBCoV4. Compounds NBCoV1 - NBCoV3 had excellent IC_50_s of 13.8-17 nM (SI: 2265-2717) in 293T/ACE2 cells, 19.3-39 nM (SI: 2282->5181) in HT1080/ACE2 cells and 98-157 nM (SI: >637->901) in A549/ACE2 cells. Compound NBCoV4 had slightly higher IC_50_s in 293T/ACE2 cells and HT1080/ACE2 cells (SI: 509 and >1852, respectively), but it displayed the 2^nd^ best activity and best SI (>1000) in A549/ACE2 cells compared to NBCoV1 - NBCoV3. NBCoV5 had poor activity against this pseudovirus in the three cell lines tested. Also, compounds, NBCoV6 - NBCoV9, displayed anti-SARS-CoV activity in all cell lines, but clearly, these compounds were less potent and displayed lower consistency than compounds at NBCoV1 - NBCoV4. In the case of SARS-CoV, the SAR followed a similar pattern to what we observed with SARS-CoV-2. This was expected since the amino acid and structural similarity of the S2 domain of the spike protein in these viruses are high.

**Table 2.** Antiviral activity of the NBCoV small molecules in a single-cycle assay in different cell lines against pseudovirus NL4-3LlEnv-Nanol uc/SARS-CoV (IC_50_ ) and selectivity index (SI ).

| Compound | 293T/ACE2 cells |  | HT1080/ACE2 cells |  | A549/ACE2 cells |  |
| --- | --- | --- | --- | --- | --- | --- |
|  | IC<sub>50</sub> (nM)<sup>a</sup> | SI | IC<sub>50</sub> (nM)<sup>a</sup> | SI | IC<sub>50</sub> (nM)<sup>a</sup> | SI |
| NBCoV1 | 17±2.6 | 2265 | 39±8.2 | 2282 | 98±4 | 878 |
| NBCoV2 | 13.8±0.2 | 2717 | 19.3±1.1 | >5181 | 111±9 | >901 |
| NBCoV3 | 17.8±4 | 2528 | 25.7±0.6 | >3891 | 157±12.5 | >637 |
| NBCoV4 | 80±2 | 509 | 54±3.5 | >1852 | 100±13 | >1000 |
| NBCoV5 | 1853±179 | 19 | 1760±330 | >57 | >2000 | N/A<sup>b</sup> |
| NBCoV6 | 175±7 | 229 | 133±9.5 | >752 | 867±62 | >115 |
| NBCoV7 | 111±1.7 | 405 | 52±8.9 | >1923 | 271±76 | >369 |
| NBCoV8 | 128±6.7 | 302 | 225±26 | >444 | 700±70 | >143 |
| NBCoV9 | 217±22 | 155 | 246±10 | >407 | 350±15 | >286 |

<sup>a</sup>The reported IC<sub>50</sub> values represent the means ± standard deviations (n = 3).
<sup>b</sup>Not Available

The NBCoV small molecules were also evaluated against MERS-CoV pseudovirus by infecting the HuH-7 cells and the MRC-5 cells. As reported in **Table 3**, we found that yet again, NBCoV1-NBCoV4 were the most potent compounds against this pseudovirus in both cell lines (IC_50:_ 95 - 158 nM in HuH-7 cells and IC_50:_ 76.5 - 123 nM in MRC-5 cells and SIs >582 and >407, respectively). NBCoV9 also had a noteworthy MERS-CoV inhibitory activity with IC_50_ of about 200 nM. NBCoV5 had no MERS-CoV inhibitory activity at the highest dose used in this assay (2000 nM), while compounds NBCoV6-NBCoV8 inhibited MERS-CoV infection with an IC_50_ lower than 1000 nM. These findings, taken together, suggest that most of the NBCoV compounds possess pancoronavirus activity. We also observed similar SAR with the MERS-CoV. Finally, to verify the specificity of the NBCoV compounds for the coronaviruses, we evaluated them against the amphotropic murine leukemia virus (A-MLV), which enters cells via macropinocytosis^52^. We found that none of the NBCoV small molecules showed appreciable activity against this control pseudovirus (IC_50_>783 nM) (**Table 4**). Based on these experiments, the data suggest that their inhibitory activity is specific to the coronaviruses. However, please note that we already mentioned earlier that NBCoV1-NBCoV9 also showed HIV-1 fusion inhibitory activity.

**Table 3.** Antiviral activity of the NBCoV small molecules in a single-cycle assay in different cell lines against pseudovirus NL4-3LlEnv-Nanoluc/MERS-CoV (IC_50_), toxicity (CC_50_), and selectivity index (SI).

| Compound | HuH-7 cells |  |  | MRC-5 cells |  |  |
| --- | --- | --- | --- | --- | --- | --- |
|  | IC<sub>50</sub> (nM)<sup>a</sup> | CC<sub>50</sub> (μM)<sup>a</sup> | SI | IC<sub>50</sub> (nM)<sup>a</sup> | CC<sub>50</sub> (μM)<sup>a</sup> | SI |
| NBCoV1 | 95±22 | ~100 | 1053 | 76.5±0.3 | 69±1 | 902 |
| NBCoV2 | 112±7 | 80±1 | 714 | 77.±0.5 | 63±2.8 | 818 |
| NBCoV3 | 158±14 | 92±2 | 582 | 123±17 | >50 | >407 |
| NBCoV4 | 131±6 | 80±6 | 611 | 80±18 | 63.5±3.5 | 794 |
| NBCoV5 | >2000 | 76±2 | N/A<sup>b</sup> | >2000 | 72.5±1 | N/A<sup>b</sup> |
| NBCoV6 | 569±31 | 71±1 | 125 | 945±77 | 71±2.8 | 75 |
| NBCoV7 | 933±153 | N/A<sup>b</sup> | N/A<sup>b</sup> | 927±63.5 | 68.8±1 | 74 |
| NBCoV8 | 509±43 | 83±2 | 163 | 559±66 | 67.5±3.5 | 121 |
| NBCoV9 | 214±23 | 55±9 | 257 | 232±17 | 42.5±3.5 | 183 |

<sup>a</sup>The reported IC<sub>50</sub> and CC<sub>50</sub> values represent the means ± standard deviations (n = 3).
<sup>b</sup>Not Available

**Table 4.** Antiviral activity of the NBCoV small molecules against control pseudovirus NL4-3.Luc.R-E-/A-MLV (IC_50_) evaluated in 293T/ACE2 cells.

| Compound | 293T-ACE2 cells/A-MLV<br>IC<sub>50</sub> (nM)<sup>a</sup> |
| --- | --- |
| NBCoV1 | 1397±12 |
| NBCoV2 | 783±76 |
| NBCoV3 | 960±57 |
| NBCoV4 | 1623±115 |
| NBCoV5 | >2000 |
| NBCoV6 | >2000 |
| NBCoV7 | >2000 |
| NBCoV8 | >2000 |
| NBCoV9 | 1627±118 |

<sup>a</sup>The reported IC<sub>50</sub> values represent the means ± standard deviations (n = 3).

### NBCoV small molecules inhibited the replication-competent authentic virus SARS-CoV- 2 (US_WA-1/2020)

As the next step, the antiviral activity of the NBCoV small molecules was evaluated by exposing Vero E6 cells with the replication-competent authentic virus SARS-CoV-2 (US_WA- 1/2020). On the third day, post-infection, the cells were observed under the microscope to evaluate the formation of virus-induced cytopathic effect (CPE). The efficacy of the small molecules was expressed as the lowest concentration capable of completely prevent virus-induced CPE (IC_100_). We found that NBCoV1 and NBCoV2 were the most efficient compounds in preventing the complete formation of CPEs with an IC_100_ of 1.25 µM followed by NBCoV3, NBCoV4, and NBCoV9, which completely prevented the formation of the virus-induced CPEs at 2.5 µM. NBCoV7 and NBCoV8 also prevented the formation of CPEs with an IC_100_ of 5 µM, while NBCoV5 and NBCoV6 did not completely prevent the formation of the virus-induced CPE at 10 µM, which was the highest dose used in this assay (**Table 5**). Moreover, when the cells were pretreated for 2 hr before infection, compounds did not show any protection against SARS-CoV-2 infection at the higher dose used in the assay (10 µM) (**Table S1**). These findings support the results obtained with the single-cycle pseudovirus-based antiviral assays.

**Table 5.** Antiviral activity (IC100) of NBCoV small molecules in Vero E6 cells infected with SARS-CoV-2 (US_WA-1/2020)

| Compound | Vero cells/<br>SARS-CoV-2 |
| --- | --- |
| | IC<sub>100</sub> (μM)<sup>a</sup> |
| NBCov1 | 1.25 |
| NBCoV2 | 1.25 |
| NBCoV3 | 2.5 |
| NBCoV4 | 2.5 |
| NBCoV5 | >10 |
| NBCoV6 | >10 |
| NBCoV7 | 5 |
| NBCoV8 | 5 |
| NBCoV9 | 2.5 |

<sup>a</sup>Values indicate the lowest concentration capable of completely preventing virus-induced CPE in 100% of the wells.

### NBCoV small molecules neutralize variants B.1.1.7 UK (Alpha), B.1.351 RSA (Beta), and B.1.617.2 India (Delta) SARS-CoV-2

Like other RNA viruses, coronaviruses depend on an error-prone RNA-dependent RNA polymerase to facilitate virus replication and adaptation ^53^. The emergence of major SARS-CoV-2 variants carrying multiple mutations in their spike is causing new concerns for increased virulence and reduced vaccine efficacy. In this study, we focused our attention on three already worldwide spread variants, the B.1.1.7 UK (Alpha), the B.1.351 RSA (Beta), and the B.1.617.2 India (Delta). We evaluated the potency of the NBCoV small molecules against these SARS-CoV-2 variants carrying single or multiple key spike mutations (B.1.1.7 UK variant: 69-70 deletion (Δ69-70)/N501Y/P681H; B.1.351 RSA: E484K/N501Y/D614G; and B.1.617.2 Delta: D614G/P681R/D950N) ^54–57^ (**Table 6**). We introduced the amino acid substitutions into the pSARS-CoV-2-S_trunc_ expression vector. Next, we infected 293T/ACE2 cells with WT and mutant SARS-CoV-2 pseudoviruses in the absence or the presence of NBCoV1 - NBCoV4 small molecules. We used NBCoV5 as a control because of its poor activity against the coronaviruses. We observed that NBCoV1 had potent antiviral activity against all mutant pseudoviruses carrying single-, double- or triple-mutations of the three variants B.1.1.7 UK, B.1.351 RSA, and B.1.617.2 Delta, as indicated by the low IC_50_s detected, which were similar to the IC_50_ obtained for the SARS-CoV-2 WT pseudovirus (**Table 6**). NBCoV2 was also a highly potent inhibitor against all mutant pseudoviruses even though the IC_50_s detected for the SARS-CoV-2 WT pseudovirus was lower than those detected against the mutant pseudoviruses (IC_50_s in the range 35-88.7 nM for all the pseudovirus variants). NBCoV4, while was highly potent against all the variants, exhibited a significant increase of the IC_50_ was against the B.1.1.7 UK triple mutant variant Δ69-70/N501Y/P681H (IC_50_ of 158 nM) and the B.1.617.2 Delta single, double and triple mutant variants (IC_50_ of 148-239 nM). We do not know precisely how these combinations of mutations cause the antiviral potency of NBCoV4 to drop; therefore, more experiments will be necessary to explain these findings. Nevertheless, the compound still retained appreciable antiviral activity against these mutants. NBCoV3 was slightly less efficient against all the mutant pseudoviruses tested, including the WT. In this case, we found a higher IC_50_ when tested against the B.1.1.7 UK triple mutant variant (IC_50_ of 232 nM). Finally, NBCoV5 had poor/no activity against these variants. These results, taken together, suggest that the NBCoV small molecules maintain their potency against the three mutant SARS-CoV-2 variants tested.

**Table 6.** Antiviral activity NBCoV small molecules against NL4-3 Env-Nanoluc/SARS-CoV-2 mutant pseudoviruses variants B.1.1.7 UK (Alpha), B.1.351 RSA (Beta), and B.1.617.2 India (Delta).

| Compound → | NBCoV1 | NBCoV2 | NBCoV3 | NBCoV4 | NBCoV5 |
| --- | --- | --- | --- | --- | --- |
| SARS-CoV-2 ↓ | IC<sub>50</sub><br>(nM)<sup>a</sup> | IC<sub>50</sub><br>(nM)<sup>a</sup> | IC<sub>50</sub><br>(nM)<sup>a</sup> | IC<sub>50</sub><br>(nM)<sup>a</sup> | IC<sub>50</sub> (nM)<sup>a</sup> |
| **WT** | 51±17 | 22.8±0.8 | 60.1±8.5 | 26±1 | 1205±240 |
| **B.1.1.7 UK (Alpha)** |  |  |  |  |  |
| N501Y | 52.5±0.2 | 49±0.5 | 44.5±0.5 | 51±0.4 | >2000 |
| Δ69-70 | 48±3 | 55±0.5 | 69±7 | 53±0.5 | >2000 |
| P681H | 35±0.5 | 51±0.3 | 55±0.3 | 44±0.4 | 1720±215 |
| N501Y/Δ69-70 | 46±0.5 | 66±1 | 94±16 | 53±11 | >2000 |
| N501Y/Δ69-70/P681H | 57±0.5 | 79±14 | 232±17 | 158±31 | >2000 |
| **B.1.351 RSA (Beta)** |  |  |  |  |  |
| E484K | 42±0.7 | 45±0.2 | 80±10.6 | 45±0.4 | >2000 |
| D614G | 56±0.6 | 61±0.3 | 145±28 | 58±0.4 | 1730±62.5 |
| N501Y/D614G | 57±2 | 59±13 | 104±1 | 77±29 | >2000 |
| E484K/N501Y | 32±0.5 | 36±0.4 | 42±1 | 51±5 | >2000 |
| E484K/D614G | 33±0.5 | 35±0.2 | 44±0.8 | 44±3 | 1846±124 |
| E484K/N501Y/D614G | 45±2 | 43±0.5 | 101±2 | 43±2.5 | >2000 |
| **B.1.617.2 India (Delta)** |  |  |  |  |  |
| D950N | 47.7±15 | 88.7±6 | N/A<sup>b</sup> | 207±29 | >2000 |
| D614G/P681R | 49±13 | 65±18 | N/Ab | 148±30 | >2000 |
| D614G/D950N | 33±10 | 26±2 | N/A<sup>b</sup> | 77.5±16.5 | >2000 |
| D614G/P681R/D950N | 58±7 | 53±12 | N/A<sup>b</sup> | 239±7 | >2000 |

<sup>a</sup>The reported IC<sub>50</sub> values represent the means ± standard deviations (n = 3).
<sup>b</sup>Not Available

### Binding affinity of the two most potent inhibitors by SPR analysis

We used Surface Plasmon Resonance (SPR) to determine the binding affinity of two of the most active inhibitors, NBCoV1 and NBCoV2, of SARS-CoV-2. We selected SARS-CoV-2 Spike (trimer) in a prefusion state as we hypothesize that these inhibitors bind to this trimer and prevent the formation of 6-HB, necessary for fusing the virus to cells. We also wanted to test any possible binding of these inhibitors to the SARS-CoV-2 spike S1 subdomains containing the RBD, which binds to the ACE2 receptor on the host cell. This method is useful in measuring the binding constant (K_D_) as well as *k*_on_ (also known as association constant, *k*a) and *k*_off_ (also known as dissociation constant, *k*_d_). Small-molecule inhibitors were passed through the chip surface, and the signal changes (in AU) of each inhibitor at varied concentrations were recorded [**Figure 3(a-d)]**. The resulting data were fit to a 1:1 binding model. The binding affinity K_D_ and kinetic parameters *k*_on_ and *k*_off_ of the target proteins’ interaction with NBCoV1 and NBCoV2 were determined (**Figure 3e**). The K_D_ value of NBCoV1 and NBCoV2 were 1.56 and 5.37 µM, respectively, with SARS-CoV-2 spike trimer. However, when these inhibitors were tested against the SARS-CoV-2 S1 subdomain, the K_D_ value of NBCoV1 was ∼5-fold higher than when bound to the prefusion S trimer. Similarly, in the case of NBCoV2, the K_D_ value was ∼9-fold higher when compared to the K_D_ value against the SARS-CoV-2 prefusion S trimer. The data indicate that these inhibitors most likely bind to the S2 subdomain of the SARS-CoV-2 trimer, although we do not know the exact mode of binding of these inhibitors. However, binding to the S2 makes sense since these inhibitors are expected to be the fusion inhibitors against coronaviruses.

**Figure 3.**
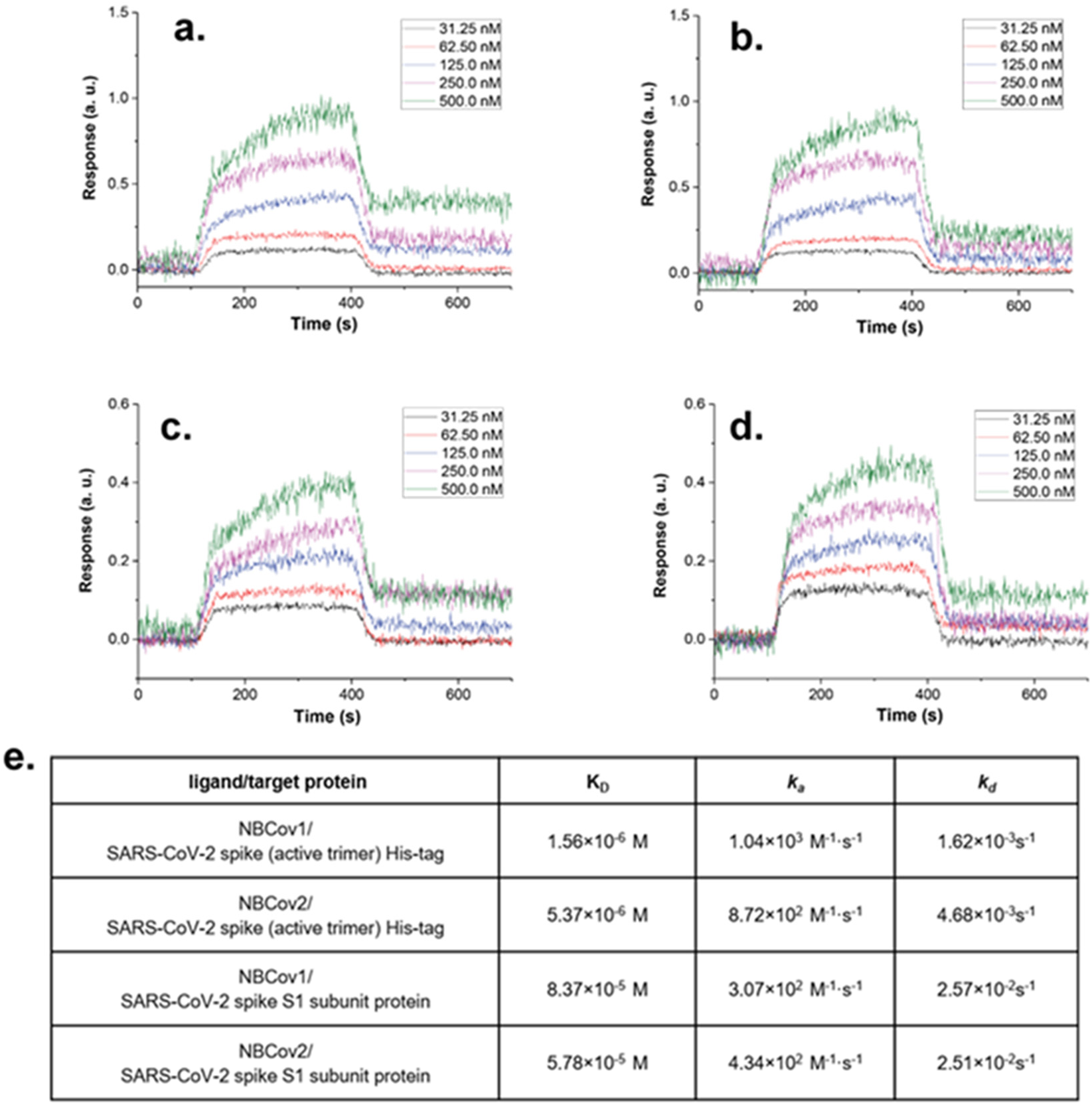
Evaluation of binding affinity of NBCoV1 and NBCoV2 to SARS-CoV -2 active trimer and SARS-CoV-2 S1 subdomain by SPR. Kinetics fitting curve (sensogram) of SARS-CoV-2 trimer to **a)** NBCoV1; **b)** NBCoV2. Kinetics fitting curve (sensogram) of SARS-CoV-2 S1 subdomain to c) NBCoV1 and **d)** NBCoV2. The binding affinity K_0_ and kinetic parameters *k_00_* and *k_0n_* of e) NBCoV1 and NBCoV2.

### NBCoV small molecules inhibited the SARS-CoV-2 mediated cell-to-cell fusion

Efficient virus spreading can be achieved by either a cell-free or a cell-associated mode involving direct cell-to-cell contact/fusion^58^. Cell-to-cell fusion mode permits the virus to infect adjacent cells without producing free virus, contributing to tissue damage and inducing syncytia formation. ACE2/SARS-CoV-2 spike interaction and subsequent conformational changes in the spike protein are critical in initiating the fusion of membranes of infected cells with the adjacent cells^59, 60^. Since we have shown that the NBCoV compounds inhibit SARS-CoV-2 and bind to the SARS-CoV-2 prefusion S trimer, we investigated whether our best compounds, NBCoV1, NBCoV2, and NBCoV4, could prevent SARS-CoV-2 mediated cell-to-cell fusion. NBCoV5 was used as a negative control, as we showed that it had no meaningful anti-SARS-CoV-2 activity.

We have set up a new and novel cell-to-cell fusion assay in our laboratory, which uses Jurkat cells expressing the luciferase gene and the SARS-CoV-2 spike wild type-WT as donor cells and 293T/ACE2 as acceptor cells. We chose Jurkat cells because they grow in suspension, and if not fused with the 293T/ACE2 cells, they can easily be removed from the wells by washing twice with PBS. Jurkat cells were pretreated with escalating concentrations of NBCoV compounds for 1 h, then added to the 293T/ACE2 cells and cocultured for 4 h to allow fusion. We found that NBCoV5 only inhibited the SARS-CoV-2 mediated cell-to-cell fusion at the higher dose (68% inhibition at 4 µM) (**Figure 4**), while NBCoV1, NBCoV2, and NBCoV4 potently inhibited the cell-to-cell fusion even at the lowest dose used in this assay. In fact, at the concentration of 250 nM, NBCoV1, NBCoV2, and NBCoV4 still maintained a 62-79% inhibitory activity of the SARS-CoV-2 mediated cell-to-cell fusion suggesting that these NBCoV small molecules interfere with the SARS-CoV-2 Spike-mediated cell-to-cell fusion.

**Figure 4.**
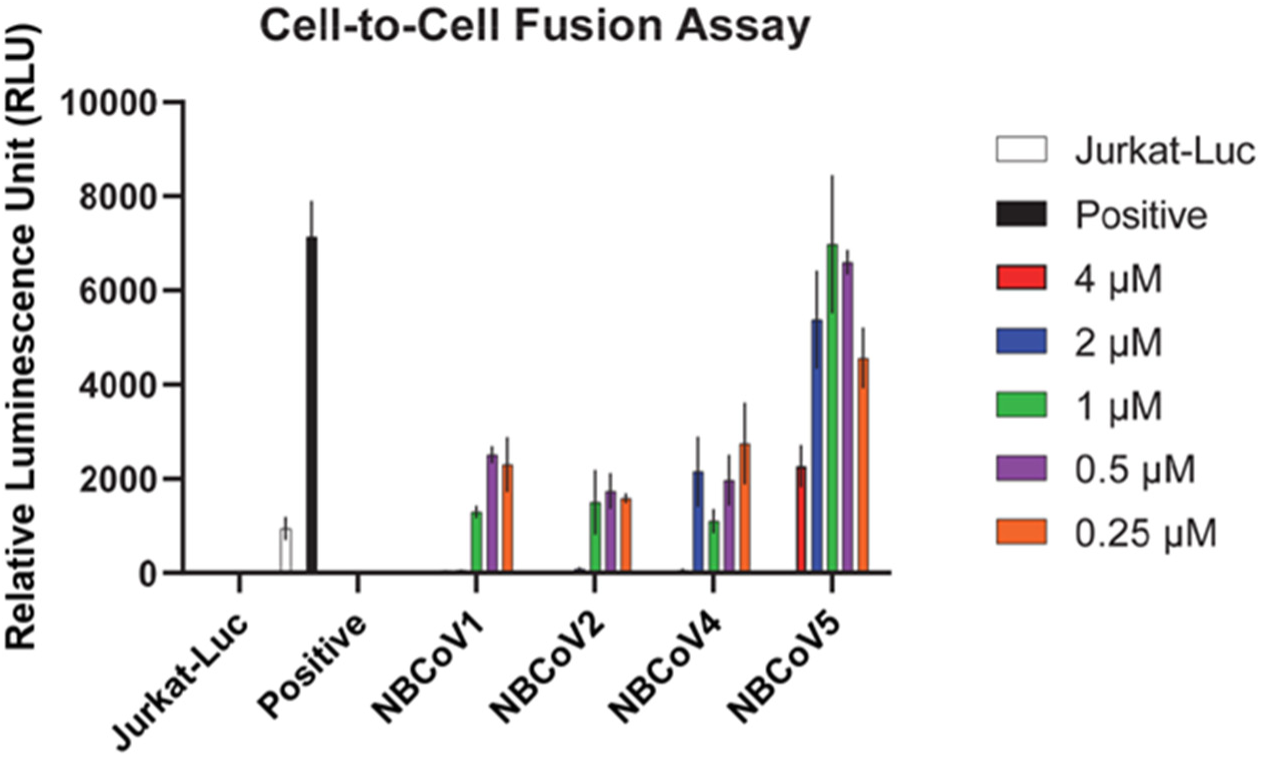
**SARS-CoV-2 mediated cell-to-cell fusion inhibition assay.** Jurkat cells expressing the SARS-CoV-2 full Spike wild-type gene from Wuhan-Hu-1 isolate and the luciferase gene were used as donor cells and the 293T/ACE2 as acceptor cells. Jurkat cells were preincubated for 1 h with different concentrations of NBCoV small molecules. Positive represent 293T/ACE2 cells cocultured with Jurkat cells in the absence of NBCoVs. Jurkat-Luc represents 293T/ACE2 cells cocultured with Jurkat cells expressing the luciferase gene only, in the absence of NBCoVs. Columns represent the means ± standard deviations (n = 2-4).

### *In vitro* ADME assessment

The *in vitro* assessment of ADME properties in the early stage of drug discovery and development, especially for the pharmaceutical industry, significantly reduced the drug attrition rate in the last two decades^61^. In 1997, the major causes of failure for drugs that advanced to clinical trials were poor ADME properties^62^. Failures in drug development during the later stages can be very costly. Therefore, we also adopted *in vitro* ADME assessments of our potent pancoronavirus inhibitors early to develop these fusion inhibitors as preclinical candidates.

We have selected one of the best inhibitors, NBCoV1, with potent antiviral activity, low cytotoxicity, and excellent SI for evaluating its ADME properties. Solubility is one of the key properties of a drug and plays a critical role in drug discovery. Therefore, we measured the solubility of the inhibitor in phosphate buffer at pK 7.4. The data in **Table 7** indicate that the solubility is NBCoV1 is low. However, there is room for further improvement through salt-formation or formulations. Due to the presence of a COOH anion in all potent NBCoV inhibitors, sodium, calcium, and potassium salts can be made to enhance the solubility and dissolution rate.^63, 64^ Next, we measured the permeability of NBCoV1 since it plays a vital role in drug absorption in the intestine and its bioavailability. Compounds with low permeability may absorb less and show poor bioavailability. The human epithelial cell line Caco-2 is the most widely used cell line to measure permeability and simulates human intestinal absorption. Therefore, we performed the Caco-2 bidirectional permeability experiment [apical to basolateral (A-B) and basolateral to apical (B-A) across the Caco-2 cell monolayer], which can be used to measure the efflux ratio and predict the human intestinal permeability of orally administered drugs. The data shown in **Table 7** indicates that the apparent permeabilities of NBCoV1 are similar to the oral drug propranolol, Papp, 10^-6^ cm/s of which was 19.7 (See *Supporting Information*). We used valspodar, a P-gp substrate, as a positive control to determine whether there was any involvement of active efflux mediated by P-gp. After treatment with 1 µM valspodar, the efflux ratio compared to no valspodar did not change, indicating that the P-gp mediated efflux was not involved.

**Table 7.**
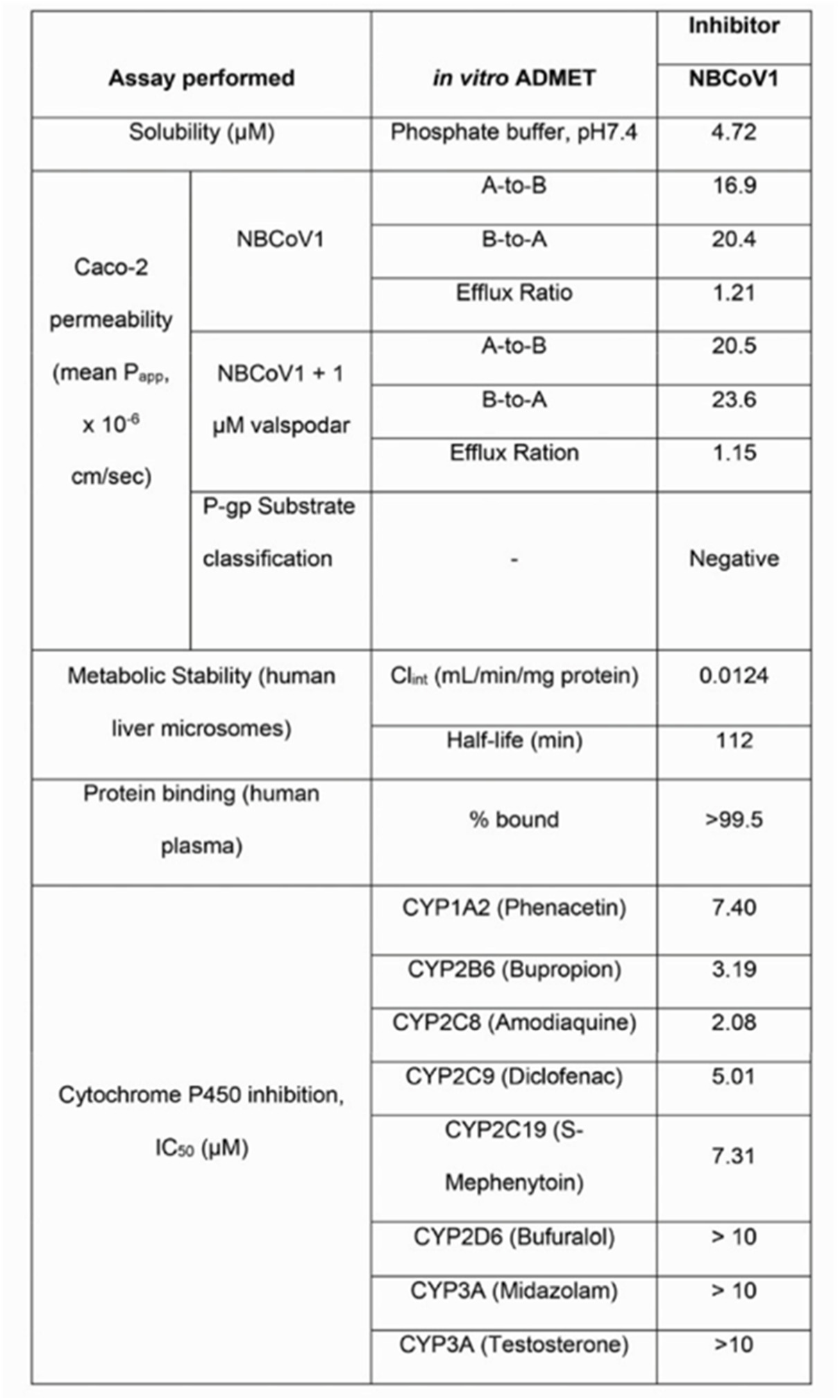
In vitro ADME profile of one of the most potent inhibitors NBCoV1

Next, we examined the metabolic stability of NBCoV1 in the human liver microsome because the liver is the most crucial site of drug metabolism in the body. The clearance data (Cl_int_) in **Table 7** indicated that NBCoV1 is a low-clearance compound, and it shows a half-life of 112 minutes. It is worthwhile to mention that achieving low clearance is often the goal of a drug discovery project to reduce drug dose, minimize exposure of the drugs in the body to reduce drug-related toxicities, and prolong half-life. Compounds with high clearance values may be cleared rapidly from the body, and the drugs may have a short duration of action and may need multiple dosing. We have also measured the binding of NBCoV1 in human plasma, and **Table 7** shows that the inhibitor is >99.5% bound. Although it may look that NBCoV1 has high protein binding, many drugs have >98% plasma protein binding, and higher protein binding does not affect the success of any drug candidates. The misconception on high plasma protein binding of drugs has been elegantly reported by Smith et al. in 2010^65^.

The cytochrome P450 (CYP450) enzyme family plays a critical role in the oxidative biotransformation of many drugs and other lipophilic xenobiotics into hydrophilic counterparts, facilitating their elimination from the body^66, 67^. There are more than 50 CYP450 enzymes in the family, but about a dozen of them, e.g., CYP1A2, CYP2B6, CYP2C8, CYP2C9, CYP2C19, CYP2D6, CYP3A4, and CYP3A5, play an essential role in metabolizing almost 80 percent of all drugs^68, 69^. Therefore, we decided to use this set of eight CYP450 enzymes to determine whether NBCoV1 has any inhibitory effects on this subfamily of enzymes that may cause potential drug-drug interactions (DDI) when co-administered with other treatment agents. DDI is a potential concern for pharmaceutical companies developing drugs and regulatory agencies such as the FDA.

- ften the following guideline is used for the CYP inhibition assessment^70^:
- IC_50_ > 10 µM (CYP inhibition low)
- < 10 µM (CYP inhibition moderate)
- < 3 µM (CYP inhibition high)

Based on the above classification, NBCoV1 showed low inhibition against CYP2D6, CYP3A, moderate inhibition against CYP1A2, CYP2B6, CYP2C9, and CYP2C19, and high inhibition against only CYP2C8 enzyme (**Table 7**). However, it is worth noting that Walsky et al. reported the inhibition of 209 drugs, and they classified high inhibition when IC_50_ < 1 µM and IC_50_ > 10 µM as moderate inhibition. In addition, this group listed felodipine, a hypertensive drug, along with five others as highly potent inhibitors^71^.

### *In vivo* pharmacokinetics (PK) of NBCoV1 and NBCoV2

Successful drug discovery depends not only on the preclinical efficacy and toxicity profile of a compound but also on the selection of the right candidate with good *in vivo* pharmacokinetics in animals (rat, dog, etc.) using appropriate dosing routes, such as oral (PO) and intravenous (IV).

We evaluated the PK parameters of two of the most active inhibitors in rats (**Table 8**) by PO and IV routes. The half-life (t_1/2_) by PO of NYBCoV1 was 11.3 hours, and IV was 3.57 hours. NBCoV1 dosed via IV showed T_max_ at 0.25 hours and PO at 2 hours, suggesting normal Clearance. The C_max_, which measures the highest drug concentration in the blood or target organ for NBCoV1 and NBCoV2, was 1499 ng/mL and 2219 ng/mL, respectively. NBCoV1 also showed an excellent mean residence time (MRT) of 14 hours. MRT measures the average time a drug molecule spends in the body and is critically important for a drug to elicit its action. The oral bioavailability of NBCoV1 was reasonably good (F%: 20) for initiating further pre-clinical studies. However, its bioavailability can be further improved through proper dosing, salt formation, or proper clinical formulation. NBCoV2 showed poor oral availability at 0.9% and half-life by PO and IV at 3.5 to 3.9 hours. NBCoV2 dosed via IV showed Cmax is at 2 hours, suggesting compound potentially precipitated after injection and redissolved to delay maximum blood levels. This suggests that the PK studies should be further evaluated by lowering the dose (1-3 mg/Kg body weight).

**Table 8.** *In vivo* PK parameters in rats of the two most active inhibitors, **NBCoV1** and **NBCoV2.**

| Parameters<sup>a</sup> | NBCoV1 |  | NBCoV2 |  | UNITS |
| --- | --- | --- | --- | --- | --- |
|  | PO | IV | PO | IV |  |
| Dose | 10 | 5 | 10 | 5 | mg/kg |
| $t_{1/2}$ | 11.32 | 3.57 | 3.96 | 3.52 | h |
| $T_{max}$ | 2 | 0.25 | 4 | 2 | h |
| $C_{max}$ | 1499.76 | 7815.76 | 30.12 | 2219.45 | ng/mL |
| $C_0$ | - | 7407.25 | - | 1710.87 | ng/mL |
| $AUC_{0-t}$ | 12023.00 | 37515.91 | 318.78 | 18074.45 | ng/mL*h |
| $MRT_{0-inf\_obs}$ | 14.34 | 4.02 | 7.28 | 5.09 | h |
| $CL_{obs}$ | - | 0.00013 | - | 0.00027 | (mg/kg)/(ng/mL)/h |
| $V_{ss\_obs}$ | - | 0.00053 | - | 0.00140 | (mg/kg)/(ng/mL) |
| $V_z/F_{obs}$ | 0.01078 | - | 0.17593 | - | (mg/kg)/(ng/mL) |
| $CL/F_{obs}$ | 0.00066 | - | 0.03082 | - | (mg/kg)/(ng/mL)/h |
| F% | 20.1 | - | 0.9 | - | 100 * AUC(PO)/AUC(IV) |

<sup>a</sup> A single oral (PO) or IV dose; $t_{1/2}$ , apparent terminal elimination half-life; $t_{max}$ , time to peak concentration; $C_{max}$ , maximum measured plasma concentration; $C_0$ , initial measured plasma concentration; AUC, area under the concentration time curve; MRT, mean residence time; CL, clearance rate of the analyte (IV only); $V_{ss}$ , volume of distribution of the analyte in the test system estimated at steady state (IV only); $V_z/F$ , apparent volume of distribution; $CL/F$ , apparent oral clearance; F%, bioavailability, represents the fraction of a dose reaching systemic circulation intact; i.e. fraction of dose absorbed.

### Do the ene-rhodanines PAINS or a part of GAINS (<u>G</u>enuinely <u>A</u>ctive <u>IN</u>hibitor<u>S</u>) in the context of pancoronavirus inhibition?

Since the publication on PAINS by Baell and Holloway^72^ and colloidal aggregators by McGovern et al. as frequent hitters or promiscuous inhibitors^73^ in high throughput screening (HTS) campaign, the awareness of these categories of compounds became part of the equation in drug discovery. However, several publications made counterarguments that all frequent hitters should not be randomly discarded without validating whether they are target-specific or true promiscuous^47, 74–77^. In a recent Editorial, Bajorath mentioned that the chemical integrity and specific biological activity of compounds containing PAINS substructure should be considered in the context of the whole compounds and how they are embedded in the structure. He also argued that“PAINS-induced activity artifacts cannot be generalized but require careful assessment on a case-by-case basis”^74^. Based on the legitimate concerns on PAINS and colloidal aggregates, nine American Chemical Society (ACS) editors outlined necessary steps to rule out any artifactual assay activity^78^. The goal of this concerted effort in authors term“not to eliminate a priori all compounds that may resemble PAINS or colloidal aggregators” but to ensure that the compounds’“behavior is well-vetted before publication.”

Ene-rhodanines have been designated as frequent hitters and speculated that the most likely activity of this series of compounds is not due to the actual target. On the contrary, Mendgen et al. in 2012 conclusively demonstrated that rhodanines and thiohydantoins possess distinct molecular interaction patterns governed by their electronic and hydrogen bonding properties and not related to promiscuous binding or aggregation. Therefore, the authors suggested not to coin these scaffolds as problematic or promiscuous binders^47^. We respect both views. Thus, in the spirit of the suggestions made by the ACS Editors^78^ and others^74, 79^, we decided to validate the antiviral activity of the set of rhodanines that we presented here.

#### a) Antiviral activity of ene-rhodanine derivatives: Is the activity due to inhibition of luciferase or direct interference with the luminescence measurement?

Initial identification of antiviral activity was performed against a lentiviral-based pseudotyped virus with spike protein from SARS-CoV-2 using a NanoLuc (luciferase-based) assay. One of the possibilities is that the ene-rhodanines used may directly inhibit the luciferase enzyme. This possibility can be ruled out as out of fourteen compounds tested, only 3-4 showed antiviral potency, although all the compounds contain the same ene-rhodanine scaffold. In neutralization assays, we pre-treated the pseudoviruses with the small molecules for 30 min before the cell infection as the target of our study is the spike protein of SARS-CoV-2. By contrast, when we pre-treated the cells rather than the viruses, we did not detect any inhibition at the higher doses used in the assay (2000 nM) (**Table S1**). These experiments suggest that 1) our compounds are not affecting the NanoLuc activity and 2) the target of these compounds is virus-related and not cell-related. In 2008, Auld et al. reported the luciferase inhibitory activity of >72,000 diverse molecules collected from a diverse chemical repository. Twenty-six rhodanines were also tested against the luciferase enzyme, and none showed any inhibitory activity^80^. In this work, to explicitly rule out that the antiviral activity of our small molecules was due to the direct inhibition of the NanoLuc and the FLuc reporters, we expressed these enzymes in 293T/17 cells. We incubated the cells lysates with 2000 nM of NBCoV small molecules (4 small molecules with the highest inhibitory activity NBCoV1-4 and two inactive compounds NBCoV5 and NBCoV34) for 10 minutes at 25 °C. As a control, lysates were untreated or treated with the Intracellular TE Nano-Glo® Substrate/Inhibitor (for the NanoLuc reporter) and 100 µM resveratrol^81^ (for the FLuc reporter). We found that while the Nanoluc inhibitor (**Figure S1a**) and the FLuc inhibitor (**Figure S1b**) completely blocked the activity of the respective luciferase enzymes, the NBCoV compounds did not affect the activity of these two enzymes (**Figure S1 a and b**). No significant differences were detected between the untreated controls and the samples treated with the NBCoV small molecules.

The second concern could be that the rhodanines may interfere with the luminescence-based measurements. However, this possibility in our case can be ruled out because we used the same luciferase enzyme in the control experiment with A-MLV-based pseudovirus as we did with cell-cell fusion assay. However, none of the rhodanine derivatives were active against A-MLV pseudovirus. On the contrary, the most active leads in the SARS-CoV-2 pseudovirus assay also showed activity against cell-cell fusion assay.

The most intriguing argument can be made with the assay results with the authentic SARS-CoV-2 virus, where no luciferase was used. The cells (Vero) were also different from those used in a pseudovirus inhibition assay. In addition, in the authentic virus assay, a microscope was used to determine the CPE, a completely different readout method. Also, in this case, when the cells (rather than the virus) were pretreated for 2 hr before infection, compounds did not confer any protection against SARS-CoV-2 infection at the higher dose used in the assay (10 µM) (**Table S1**). All these concurrent assays firmly establish that the target of the rhodanines is not random but more specific to the spike protein of SARS-CoV-2.

#### b) Target specificity measured by a direct binding study by SPR

As suggested by the ACS Editors^78^, we went a step further by measuring the direct binding of the most active inhibitors to the target SARS-CoV-2 spike protein by surface plasmon resonance (SPR) method. Since we hypothesized that these inhibitors are expected to bind the HR1 domain of the spike protein and prevent the six-helix bundle formation similar to HIV-1 or other Class I fusion proteins of enveloped viruses, we used a SARS-CoV-2 prefusion S trimer. Both NBCoV1 and NBCoV2 showed low µM K_D_. Thus, although these inhibitors showed some binding to the S1 domain, the K_D_ values were 5-9-fold higher. Therefore, this critical study demonstrated that these inhibitors were preferentially bound to the Spike protein’s prefusion state and supported our hypothesis. Admittedly, we do not know the exact binding location, but future x-ray or Cryo-EM structure determination with the inhibitors will undoubtedly provide us with a wealth of information.

#### c) Do the inhibitors bind specifically to the virus spike protein or promiscuously to some cellular proteins?

In 2014, Baell and Walters, in their comments published in Nature, coined ene-rhodanines as one of the“most insidious” offenders^82^. They also mentioned that this scaffold primarily works through covalent modification and metal complexation. We do not know whether our inhibitors participate through any such mechanisms without any structural information. However, we will present some experimental evidence to show that the inhibitors bind to the virus part, not the cellular component, most likely not through the ene-rhodanine scaffold. To accomplish our goal, we used two-prong approaches.

##### 1) The time of addition of compounds is critical for their antiviral activity

Pseudovirus-base inhibition assay: Entry/fusion inhibitors that target the virus envelope/spike proteins do not show inhibition if added to the cell first, followed by virus addition. We successfully demonstrated that when we incubated compounds with the cells first and then added pseudovirus, none of the compounds showed any inhibition (**Table S1**). However, if we reverse the sequence by pre-incubating the virus with the compound before adding it to the cells, compounds show dose-response inhibition (**Tables 1, 2, 3 and 6**).
Authentic live SARS-CoV-2 virus: We demonstrated a similar effect with live virus assay (**Table 5**). None of the compounds tested showed any inhibition when first added to the cells (**Table S1**).

The above experiments conclusively established that the target of the inhibitors was not the cellular components but the virus itself.

##### 2) Cellular toxicity vs. antiviral activity

During pseudovirus-based inhibition assay, cellular toxicity was also assessed for each compound without adding any virus. The data in **Table 1** and **Table 3** indicate all active compounds have low cytotoxicity making the SI values anywhere from >586 to 4000. If the compounds target the cellular component, then the SI values would have been much lower. Based on the ACS Editors’ recommendation, we demonstrated that“the compound is active at a concentration substantially lower than those producing cellular toxicity”^78^.

#### d) Are the NBCoV small molecules promiscuous aggregation-based inhibitors?

Based on the steps outlined by the nine American Chemical Society (ACS) editors to rule out inhibitory activity due to colloidal aggregation^78^, we decided to investigate these compounds further. Based on one of the suggestions by the Editors to use the publicly available filters, we used the online software, Advisor, developed by Shoichet’s team at UCSF (http://advisor.bkslab.org/). The software returned with a message that none of the compounds were like any known aggregator in their database (Supporting Information, **Figure S2**). However, it also alerted that since the molecules are hydrophobic, other appropriate tests should be performed. The authors suggested that if the activity of the aggregation-based inhibitor can be attenuated by small concentrations of nonionic detergent (0.025% Tween-80), the compound is likely an aggregator ^83, 84^. Also, a colloidal aggregator mostly exhibits a steep dose-response curve, and it may be precipitated by centrifugation ^85–88^. Due to the different sensitivity of the cells to detergents, we initially performed a cytotoxicity assay with 293T/ACE2 cells in the presence and absence of 0.025% of Tween-80. Unfortunately, we found that even such a low concentration of Tween-80 was inducing significant cytotoxicity (**Figure S3a**). Moreover, in the presence of 0.025% Tween-80, the infection of 293T/ACE2 cells with SARS-CoV2 pseudovirus was dramatically decreased (**Figure S3b**) compared to the infection done in the absence of Tween-80, suggesting that 0.025% of Tween-80 may also be affecting the viral viability along with the cell viability. Then, we decided to perform the neutralization assay using the supernatant of aliquots of NBCoV1, which were centrifuged to eliminate ‘eventual’ colloidal aggregations. The dose-response obtained with the centrifuged NBCoV1 was very similar to that obtained with the not-centrifuged NBCoV1 (NBCoV1-control) (**Figure S3c**). There was no difference in the calculated IC_50_s, which were 50 nM for the NBCoV1-control and 45 nM for the NBCoV1-centrifuged, suggesting that NBCoV1 activity was not due to colloidal aggregation.

Additionally, in 2003 Seidler et al. ^89^ suggested that potential aggregators can be screened for inhibition of three unrelated enzymes, specifically, ß-lactamase, trypsin, and malate dehydrogenase (MDH), which are highly sensitive to compound aggregation. One of their criteria suggested that it can be considered promiscuous if the compound inhibits all three enzymes. To this end, we evaluated the activity of 6 NBCoV compounds (4 compounds with the highest inhibitory activity NBCoV1-4 and two inactive compounds NBCoV5 and NBCoV34) at 2000 nM against those three enzymes using a colorimetric assay. As shown in **Table S2**, we found that the NBCoV compounds had no inhibitory activity against the three enzymes ß-lactamase, trypsin, and MDH, indicating that these compounds should not be considered further as aggregators.

Furthermore, as per the ACS editors recommendation^78^, we demonstrated through high SI values that antiviral activity of the most potent inhibitors is due to actual inhibition of the virus infection to cells, not due to cellular toxicity.

Therefore, we demonstrated through a series of rationale and control experiments as per the ACS Editors’ and others’ ^78^ recommendations that the pancoronavirus inhibitors presented in this article genuinely target the viral component, specifically the spike protein, and elicit true antiviral potency.

## CONCLUSIONS

Based on the remarkable similarity in the mechanism of fusion of coronaviruses spike protein and envelope glycoproteins of HIV-1, we have discovered a series of pancoronavirus fusion inhibitors, which also show potent inhibition against the COVID-19 variants recently identified in the UK (Alpha), South Africa (Beta), and India (Delta). Out of thirteen compounds tested, we found at least three of them showed low nM IC_50_ in a pseudovirus-based inhibition assay. These molecules also showed complete inhibition of CPE (IC_100_) against an authentic live virus, SARS-CoV-2 (US_WA-1/2020), tested in Vero cells. Although limited, the SAR indicates that a balance of electrostatic and hydrophobic interactions is needed for optimum antiviral activity. For example, when phenylethyl moiety was replaced by H or smaller hydrophobic groups, the inhibitory activity of those compounds disappeared. The SAR also shows that there is room for further derivatization of the phenylethyl moiety. The direct binding study by SPR confirmed that these molecules bind to the prefusion trimer of the spike protein of SARS-CoV-2 more tightly than the S1 subdomain of the spike protein. Subsequent cell-to-cell fusion assay confirmed that these inhibitors efficiently prevent virus-mediated cell-to-cell fusion. We also demonstrated through a series of rationally designed experiments that these inhibitors are not promiscuous but true pancoronavirus inhibitors despite the presence of ene-rhodanine scaffold, which was termed by some as “frequent hitters”. As part of our early drug discovery protocol, we also performed the ADME study. It indicated that the solubility of these inhibitors needs further improvement either through chemical modifications or through salt formation. All other ADME properties measured showed drug-like characteristics. Furthermore, the pharmacokinetic (PK) study in rats demonstrated that NBCoV1 has all the desirable features, including 20% oral availability to be considered for further pre-clinical assessments. Overall, we discovered a set of novel small-molecule pancoronavirus fusion inhibitors, which are likely candidates with great potential to be developed as therapy of COVID-19 and related coronavirus diseases.

## EXPERIMENTAL SECTION

### Cells and plasmids

The MRC-5, A549, HT-1080, HeLa, HEK293T, and HEK293T/17 cells were purchased from ATCC (<u>Manassas, VA</u>). The Human Lung carcinoma (A549) cells expressing Human Angiotensin-Converting Enzyme 2 (HA FLAG) (Catalog No. NR-53522) were obtained from BEI Resources, NIAID, NIH. The Human T-Cell Lymphoma Jurkat (E6-1) cells were obtained through the NIH ARP. The HuH-7 (JCRB0403) cells were obtained from JCRB Cell Bank (Osaka, Japan). The HT1080/ACE2 (human fibrosarcoma) cells, the 293T/ACE2 cells, and the two plasmids pNL4-3ΔEnv-NanoLuc and pSARS-CoV-2-S_Δ19_ were kindly provided by Dr. P.Bieniasz of Rockefeller University^48^. The pSV-A-MLV-Env (envelope) expression vector ^90, 91^ and the Env-deleted proviral backbone plasmids pNL4-3.Luc.R-E-DNA ^92, 93^ were obtained through the NIH ARP. The two plasmids pSARS-CoV and pMERS-Cov were kindly provided by Dr. L. Du of New York Blood Center. The expression vector containing SARS-CoV-2 full Spike wild-type (WT) gene from Wuhan-Hu-1 isolate (pUNO1-SARS-S) was purchased from InvivoGen (San Diego, CA). The pFB-Luc plasmid vector was purchased from Agilent Technologies (Santa Clara, CA).

### Small Molecules

We screened a set of nine 3-(5-((4-oxo-3-phenethyl-2-thioxothiazolidin-5-ylidene)methyl)furan-2-yl)benzoic acids from our stock (NYBC). The details of the synthesis, purification and analytical characterization were published earlier^42^. We also purchased NBCoV1 from Sigma-Aldrich (St. Louis, MO) in larger quantities. This compound was purified and characterized thoroughly by our group (compounds purity is >95%) - details are in the Supporting Information). We also purchased one control analog without the COOH group, 5- ((5-(4-chlorophenyl)furan-2-yl)methylene)-3-phenethyl-2-thioxothiazolidin-4-one (NBCoV15), and NBCoV17, NBCoV28 and NBCoV34 from Chembridge Corporation (San Diego, CA). (All purchased compounds purity is >95% -details of the analyses are reported in the Supporting Information)

### Pseudoviruses preparation

To prepare pseudoviruses capable of single-cycle infection, 8×10^6^ HEK293T/17 cells were transfected with a proviral backbone plasmid and an envelope expression vector by using FuGENE HD (Promega, <u>Madison, WI</u>) and following the manufacturer’s instructions. To obtain the SARS-CoV-2, SARS-CoV and the MERS-CoV pseudoviruses, the cells were transfected with the HIV-1 Env-deleted proviral backbone plasmid pNL4-3ΔEnv-NanoLuc DNA and the pSARS-CoV-2-S_Δ19_^48^, pSARS-CoV and pMERS-CoV Env plasmids, respectively. For the A-MLV pseudovirus, the cells were transfected with the Env-deleted proviral backbone plasmids pNL4-3.Luc.R-.E-DNA and the pSV-A-MLV-Env expression vector. Pseudovirus-containing supernatants were collected two days after transfection, filtered, tittered, and stored in aliquots at −80 °C. Pseudovirus titers were determined to identify the 50% tissue culture infectious dose (TCID_50_) by infecting the different cell types. For the titers in HT1080/ACE2 cells, 2×10^4^ cells were added to 100-μL aliquots of serial 2-fold dilutions of pseudoviruses in a 96-well plate and incubated for 24 h. For the titers in A549/ACE2 cells, 1×10^4^ cells were added to 100-μL aliquots of serial 2-fold dilutions of pseudoviruses in a 96-well plate and incubated for 48 h. For the titers in 293T/ACE2, MRC-5, and HuH-7 cells, 1×10^4^ cells/well were plated in a 96-well plate and incubated overnight before adding the 100-μL aliquots of serial 2-fold dilutions of pseudoviruses and incubated for 48h. Following the incubation time, the cells were washed with PBS and lysed with 50 μL of the cell culture lysis reagent (Promega, <u>Madison, WI</u>). For the SARS-CoV-2 titers, 25 µL of the lysates were transferred to a white plate and mixed with the same volume of Nano-Glo® Luciferase reagent (Promega). For the A-MLV titers, 25 µL of the lysates were transferred to a white plate and mixed with 50µL of luciferase assay reagent (Luciferase assay system, Promega). We immediately measured the luciferase activity with a Tecan SPARK multifunctional microplate reader (Tecan, Research Triangle Park, NC). The wells producing relative luminescence unit (RLU) levels 10 times the cell background was scored as positive. We calculated the TCID_50_ according to the Spearman-Karber method^94^.

### Analysis of the incorporation of the spike proteins into SARS-CoV-2, SARS-CoV, and MERS-CoV pseudoviruses

To confirm the incorporation of the respective spike proteins into the SARS-CoV-2, SARS-CoV, and MERS-CoV pseudoviruses, 2 mL of the pseudovirus-containing supernatants were ultra-centrifuged for 2 h at 40,000 rpm on a 20 % sucrose cushion to concentrate the viral particles. Viral pellets were lysed and processed for protein analysis. The viral proteins were resolved on a NuPAGE Novex 4–12 % Bis-Tris Gel (Invitrogen, <u>Carlsbad, CA</u>). The SARS-CoV-2 and SARS-CoV viral lysates were immuno-detected with a SARS spike protein antibody (NB-100-56578, Novus Biological, <u>Littleton, CO</u>), followed by an anti-rabbit-IgG HRP linked whole antibody (GE Healthcare, <u>Chicago, IL</u>). The MERS-CoV viral lysate was immuno-detected with a MERS-coronavirus spike protein S2 polyclonal antibody (Invitrogen, Carlsbad, CA) followed by a donkey anti-rabbit IgG (H+L), HRP secondary antibody. Proteins were visualized using chemiluminescence.

### Evaluation of the ACE2 and CD26 (DPP4) expression

The expression of the ACE2 receptor and the DDP4 receptor in the different cell lines was evaluated by Western Blot to find a correlation with the infection levels detected in the different cell lines (HT-1080\ACE2 and HT-1080, A549\ACE2, A549, 293T/ACE2, HEK293T and HeLa for SARS-CoV-2 and SARS-CoV and HuH-7, MRC-5 and HeLa for MERS-CoV). Cell pellets were lysed and processed for protein analysis. For Blot 1 we loaded 50 µg of proteins (293T/ACE2, HEK293T, HT1080/ACE2, HT1080, and HeLa), and for Blot 2, we loaded 75 µg of proteins (A549/ACE2, A549, and HeLa). For Blot 3 we loaded 50 µg of proteins (MRC-5, HuH-7, and HeLa). The proteins were resolved on a NuPAGE Novex 4–12 % Bis-Tris Gel. Blot 1 and Blot 2 were immuno-detected with a human anti-ACE2 mAb (AC384) (Adipogen Life Sciences, San Diego, CA). The ECL Mouse IgG, HRP-linked whole Ab (from sheep) (Amersham, Little Chalfont, UK) was used as a secondary antibody. Blot 3 was immuno-detected with a human DPP4 Monoclonal Antibody (OTI11D7), TrueMAB™ (Invitrogen), followed by the ECL Mouse IgG, HRP secondary antibody. Cell lysates were also immuno-detected with the housekeeping gene β-actin as a loading control. Proteins were visualized using chemiluminescence.

Additionally, the correlation of the pseudovirus SARS-CoV-2 and SARS-CoV with the expression of the ACE2 receptor was analyzed by infecting cells expressing different amounts of the ACE2 receptor with the same volume of the pseudovirus-containing supernatant. Briefly, 50 µL of SARS-CoV-2 and SARS-CoV diluted with 50 µL serum-free medium was added to wells of a 96-well cell culture plate. Next, the cells were added as follow: HT-1080\ACE2 and HT-1080 cells were added to the respective wells at the concentration of 2×10^4^ cells/well and incubated for 24 h at 37°C; A549\ACE2, A549, and HeLa cells were added to the respective wells at a concentration of 1×10^4^ cells**/**well and incubated for 48 h at 37°C. For the 293T/ACE2 and 293T, 1×10^4^ cell/well were plated the day before, then infected with the same volume of SARS-CoV-2 and SARS-CoV. Uninfected cells for all cell lines were used as a negative control.

The correlation of infection of the MERS-CoV pseudovirus with the expression of the CD26 (DPP4) receptor was analyzed by infecting three different cell types (MRC-5, HuH-7, and HeLa cells) with the same volume of the MERS-CoV pseudovirus-containing supernatant. Uninfected cells for all cell lines were used as a negative control. Following the incubation time, the cells were washed with PBS and lysed with 50 μL of the cell culture lysis reagent. Twenty-five µL of the lysates were transferred to a white plate and mixed with the same volume of Nano-Glo® Luciferase reagent. The luciferase activity was immediately measured with a Tecan SPARK. **Measurement of antiviral activity**

The antiviral activity of the NBCoV small molecules was evaluated in a single-cycle infection assay by infecting different cell types with the SARS-CoV-2, SARS-CoV, or MERS-CoV pseudoviruses as previously described with minor modifications ^27, 49^. **HT1080/ACE2 cells.** Briefly, in 96-well culture plates, aliquots of SARS-CoV-2 or SARS-CoV at about 3000 TCID_50_/well at a multiplicity of infection (MOI) of 0.1 were pre-incubated with escalating concentrations of the NBCoV small molecules for 30 min. Next, 2×10^4^ cells were added to each well and incubated at 37°C. HT1080/ACE2 cells cultured with medium with or without the SARS-CoV-2 or SARS-CoV pseudoviruses were included as positive and negative controls, respectively. Following 24 h incubation, the cells were washed with 200 µL of PBS and lysed with 50 µL of lysis buffer. 25 µL of the lysates were transferred to a white plate and mixed with the same volume of Nano-Glo® Luciferase reagent. The luciferase activity was measured immediately with the Tecan SPARK. The percent inhibition by the small molecules and the IC_50_ (the half-maximal inhibitory concentration) values were calculated using the GraphPad Prism 9.0 software (San Diego, CA).

#### A549/ACE2 cells

For the evaluation of the antiviral activity in A549/ACE2 cells, aliquots of the pseudovirus SARS-CoV-2 or SARS-CoV at about 1500 TCID_50_/well at an MOI of 0.1 were pre-incubated with escalating concentrations of the NBCoV small molecules for 30 min. Next, 1×10^4^ cells were added to each well and incubated. A549/ACE2 cells cultured with medium with or without the SARS-CoV-2 or SARS-CoV pseudoviruses were included as positive and negative controls, respectively. Following 48 h incubation, the cells were washed with PBS and lysed with 50 µL of lysis buffer. Twenty-five µL of the cell lysates were processed as reported above to measure the luciferase activity and calculate the percent inhibition by the NBCoV small molecules and the IC_50_.

#### 293T/ACE2 cells

We evaluated the antiviral activity of NBCoV small molecules in 293T/ACE2 cells infected with pseudoviruses SARS-CoV-2 and SARS-CoV. Briefly, 96-well plates were coated with 50 µL of poly-l-Lysine (Sigma-Aldrich, St. Louis, MO) at 50 µg/mL. Following 3 h incubation at 37°C, the plates were washed with PBS and let to dry. The 293T/ACE2 cells were then plated at 1×10^4^/well and incubated overnight. On the following day, the aliquots of the pseudoviruses at about 1500 TCID_50_/well at an MOI of 0.1 were pretreated with graded concentrations of the NBCoV small molecules for 30 min and added to the cells. 293T/ACE2 cells cultured with medium with or without the SARS-CoV-2 or SARS-CoV pseudoviruses were included as positive and negative controls, respectively. Additional experiments were performed by pre-treating the cells rather than the virus, with escalating concentration of NBCoV small molecules for 30 min before infection with pseudoviruses SARS-CoV-2. After 48 h incubation, the cells were washed with PBS and lysed with 50 μL of lysis buffer. Twenty-five µL of the cell lysates were processed as reported above to measure the luciferase activity and calculate the percent inhibition by the NBCoV small molecules and the IC_50_. Additionally, to test the specificity of the small molecules, we evaluated their activity against pseudovirus A-MLV at about 1500 TCID_50_/well at an MOI of 0.1 by following the infection protocol described above. Following 48 h incubation, 25 µL of the lysates were transferred to a white plate and mixed with 50 µL of a luciferase assay reagent. The luciferase activity was immediately measured.

#### MRC-5 and HuH-7 cells

For the neutralization assay in MRC-5 and HuH-7 cells, 1×10^4^ cells/well were plated in a 96-well cell culture plate and incubated overnight. On the following day, aliquots of the MERS-CoV pseudovirus at about 1500 TCID_50_/well at an MOI of 0.1 were pretreated with graded concentrations of the small molecules for 30 min and added to the cells. MRC-5 and HuH-7 cells cultured with medium with or without the SARS-CoV-2 or SARS-CoV pseudoviruses were included as positive and negative controls, respectively. After 48 h incubation, the cells were washed and lysed with 50 μL of lysis buffer (Promega). Twenty-five µLs of the cell lysates were processed as reported above to measure the luciferase activity and calculate the percent inhibition by the NBCoV small molecules and the IC_50_.

### SARS-CoV-2 Microneutralization Assay

The standard live virus-based microneutralization (MN) assay was used ^95–97^. Briefly, serially two-fold and duplicate dilutions of individual NBCoV small molecules were incubated with 120 *plaque-forming unit* (*PFU*) of SARS-CoV-2 (US_WA-1/2020) at room temperature for 1 h before transferring into designated wells of confluent Vero E6 cells (ATCC, CRL-1586) grown in 96-well cell culture plates. Vero E6 cells cultured with medium with or without the same amount of virus were included as positive and negative controls, respectively. Additional experiments were performed by pre-treating the Vero cells, with an escalating concentration of NBCoV small molecules for 2 h before infection with SARS-CoV-2. After incubation at 37°C for 3 days, individual wells were observed under the microscope to determine the virus-induced formation of cytopathic effect (CPE). The efficacy of individual drugs was expressed as the lowest concentration capable of completely preventing virus-induced CPE in 100% of the wells.

### Evaluation of cytotoxicity

The evaluation of the cytotoxicity of NBCoV small molecules in the different cell types was performed in parallel with the antiviral activity assay and measured by using the colorimetric CellTiter 96® AQueous One Solution Cell Proliferation Assay (MTS) (Promega, <u>Madison, WI</u>) following the manufacturer’s instructions.

#### HT1080/ACE2 cells

Briefly, aliquots of 100 µL of the NBCoV small molecules at graded concentrations were incubated with 2×10^4^/well HT1080/ACE2 cells and cultured at 37 °C. Following 24 h incubation, the MTS reagent was added to the cells and incubated for 4 h at 37 °C. The absorbance was recorded at 490 nm. The percent of cytotoxicity and the CC_50_ (the concentration for 50 % cytotoxicity) values were calculated as above.

#### A549/ACE2 cells

For the cytotoxicity assay in A549/ACE2 cells, aliquots of escalating concentrations of the small molecules were incubated with 1×10^4^/well A549/ACE2 cells and cultured at 37 °C. Following 48 h incubation, the MTS reagent was added to the cells and incubated for 4 h at 37 °C. The absorbance was recorded, and the percent of cytotoxicity and the CC_50_ values were calculated as above.

#### HuH-7, MRC-5 and 293T/ACE2 cells

For the cytotoxicity assay in HuH-7, MRC-5, and 293T/ACE2 cells, 1×10^4^/well cells were plated in a 96-well cell culture plate and incubated overnight. The following day, aliquots of escalating concentrations of the NBCoV compounds were added to the cells and incubated at 37 °C. Following 48 h incubation, the MTS reagent was added to the cells and incubated for 4 h at 37 °C. The absorbance was recorded at 490 nm. The percent of cytotoxicity and the CC50 values were calculated as above.

### Drug sensitivity of spike-mutated pseudovirus

Amino acid substitutions or deletions were introduced into the pSARS-CoV-2-S_trunc_ expression vector by site-directed mutagenesis (Stratagene, La Jolla, CA) using mutagenic oligonucleotides as follow:

SaCoV2-E484K-F: ACCCCTTGTAACGGCGTG**A**AAGGCTTCAACTGCTACTTCCCA

SaCoV2-E484K-REV: TGGGAAGTAGCAGTTGAAGCCTT**T**CACGCCGTTACAAGGGG

SaCoV2-N501Y-F: TCCTACGGCTTTCAGCCCACA**T**ATGGCGTGGGCTATCAGCCC

SaCoV2-N501Y-REV: GGGCTGATAGCCCACGCCAT**A**TGTGGGCTGAAAGCCGTAGGA

SaCoV2-D614G-F: CAGGTGGCAGTGCTGTACCAGG**G**CGTGAACTGTACCGAAGTG

SaCoV2-D614G-REV: CACTTCGGTACAGTTCACG**C**CCTGGTACAGCACTGCCACCTG

SaCoV2-P681H-F: CAGACACAGACAAACAGCC**A**CAGACGGGCCAGATCTGTG

SaCoV2-P681H-REV: CACAGATCTGGCCCGTCTG**T**GGCTGTTTGTCTGTGTCTG

SaCoV2-P681R-F: CAGACACAGACAAACAGCC**G**CAGACGGGCCAGA TCTGTG

SaCoV2-P681R-REV: CACAGATCTGGCCCGTCTG**C**GGCTGTTTGTCTGTGTCTG

SaCoV2-D950N-F: GCCCTGGGAAAGCTGCAG**A**ACGTGGTCAACCAGAATGCC

SaCoV2-D950N-REV: GGCATTCTGGTTGACCACGT**T**CTGCAGCTTTCCCAGGGC

SaCoV2-Δ(69-70)-S: GTGACCTGGTTCCACGCCATCTCCGGCACCAATGGCA CCAAG

SaCoV2-Δ(69-70)-REV: CTTGGTGCCATTGGTGCCGGAGATGGCGTGGAACCAGGTCAC

and following the manufacturer’s instructions. Site mutations were verified by sequencing the entire spike gene of each construct. To obtain the SARS-CoV-2 pseudovirus carrying the amino acid substitutions, the cells were transfected with the HIV-1 Env-deleted proviral backbone plasmid pNL4-3ΔEnv-NanoLuc DNA and the mutant pSARS-CoV-2-S_Δ19_ as described above. Pseudoviruses were tittered by infecting 293T/ACE2 cells as described above. To measure the activity of the compounds against the pseudoviruses expressing different point mutations, 293T/ACE2 cells were infected with the ENV-mutated pseudoviruses pretreated for 30 min with different concentrations of the NBCoV compounds and incubated for 2 days, as described above. Cells were washed with PBS and lysed with 50 μL of cell culture lysis reagent. Twenty-five µL of the cell lysates were processed as reported above to measure the luciferase activity and calculate the percent inhibition and the IC_50_.

### Cell-to-Cell fusion inhibition assay

For the SARS-CoV-2 mediated cell-to-cell fusion assay, we used Jurkat cells which transiently expressed the luciferase gene and stably expressed the SARS-CoV-2 full Spike wild-type (WT) gene from Wuhan-Hu-1 isolate as donor cells and the 293T/ACE2 as acceptor cells. Briefly, Jurkat cells at 2×10^5^/mL were transfected with 1 µg/mL of SARS-CoV-2 WT expression vector by using 5 µL/mL of FuGene HD and following the manufacturer’s instructions. Following 24 h incubation, transfected Jurkat cells were washed and selected for the SARS-CoV-2 spike expression using Blasticidin at a concentration of 10 µg/mL. To rule out cell resistance to the antibiotic, Jurkat cells that were not transfected with the SARS-CoV-2 spike were exposed to the same concentration of Blasticidin in parallel; the culture was depleted entirely in about 14 days. The antibiotic was replaced every four days, and the selection lasted for about 20 days. On the day before the assay, the 293T/ACE2 were plated in a 96-well cell culture plate at 8×10^4^/well, while the Jurkat cells were washed with PBS to remove the Blasticidin, resuspended at 2×10^5^/mL, and transfected with 1 µg/mL of pFB-Luc expression plasmid DNA using 5 µL/mL of FuGene HD. Following 20 h incubation, the Jurkat cells were washed with PBS, and aliquots of 8×10^4^/well were incubated with escalating concentrations of the NBCoV compounds for 1 h. Finally, the Jurkat cells were transferred to the respective wells containing the 293T/ACE2 cells. Additionally, 293T/ACE2 cells cultured with medium with or without the Jurkat cells were included as positive and negative controls, respectively. As an additional control, a set of 293T/ACE2 cells were incubated with Jurkat cells expressing the luciferase gene only (Jurkat-Luc). The plate was spun for 5 minutes at 1500 rpm then incubated for 4 h at 37°C. The wells were carefully washed twice with 200 µL of PBS to remove the Jurkat cells that did not fuse with the 293T/ACE2 cells. Finally, the cells were lysed to immediately measure the luciferase activity to calculate the percentage of inhibition of the SARS-CoV-2 mediated cell-to-cell fusion.

### Binding analysis by SPR

The binding study of two of the most active small-molecule inhibitors was performed by Profacgen, New York, NY. The bare gold-coated (thickness 47 nm) PlexArray Nanocapture Sensor Chip (Plexera Bioscience, Seattle, WA, US) was prewashed with 10× PBST for 10 min, 1× PBST for 10 min, and deionized water twice for 10 min before being dried under a stream of nitrogen prior to use. Various concentrations of biotinylated proteins dissolved in water were manually printed onto the Chip with Biodo bioprinting at 40% humidity via biotin-avidin conjugation. Each concentration was printed in replicate, and each spot contained 0.2 μL of sample solution. The chip was incubated in 80% humidity at 4°C overnight and rinsed with 10× PBST for 10 min, 1× PBST for 10 min, and deionized water twice for 10 min. The chip was then blocked with 5% (w/v) non-fat milk in water overnight and washed with 10× PBST for 10 min, 1× PBST for 10 min, and deionized water twice for 10 min before being dried under a stream of nitrogen prior to use. SPRi measurements were performed with PlexAray HT (Plexera Bioscience, Seattle, WA, US). Collimated light (660 nm) passes through the coupling prism, reflects off the SPR-active gold surface, and is received by the CCD camera. Buffers and samples were injected by a non-pulsatile piston pump into the 30 μL flowcell that was mounted on the coupling prism. Each measurement cycle contained four steps: washing with PBST running buffer at a constant rate of 2 μL/s to obtain a stable baseline, sample injection at 5 μL/s for binding, surface washing with PBST at 2 μL/s for 300 s, and regeneration with 0.5% (v/v) H3PO4 at 2 μL/s for 300 s. All the measurements were performed at 25°C. The signal changes after binding and washing (in AU) are recorded as the assay value.

Selected protein-grafted regions in the SPR images were analyzed, and the average reflectivity variations of the chosen areas were plotted as a function of time. Real-time binding signals were recorded and analyzed by Data Analysis Module (DAM, Plexera Bioscience, Seattle, WA, US). Kinetic analysis was performed using BIAevaluation 4.1 software (Biacore, Inc.).

#### *In vitro* ADME Study

Details of the *in vitro* ADME study and data analyses can be found in the *Supporting Information*.

#### *In vivo* pharmacokinetics in rats

We selected two of the most active inhibitors, NBCoV1 and NBCoV2, to evaluate the pharmacokinetics (PK) in rats. Rats were 11 weeks and 1 day old and weighed between 200-250 grams. A total of twelve (12) Female Sprague Dawley (SD) rats (Charles River Laboratory, USA) were implanted with a jugular vein catheter and were assigned to the study following acclimation for seven (7) days. Rats were divided into four (4) treatment groups consisting of three (3) rats each. On Day 0, 10mg/kg/animal was administered via oral gavage for groups 1 and 3. On Day 0, 5 mg/kg/animal was administered via tail vein injection based on their body weights for groups 2 and 4. All animals underwent blood collection for plasma at 5 minutes, 15 minutes, 30 minutes, 1 hour, 2 hours, 4 hours, 8 hours, and 24 hours post-dosing. At 24-hour post-dosing, all animals were euthanized post terminal blood collection without performing a necropsy. The study was conducted under BSL-1 safety conditions.

The concentrations of the Test Article in plasma were determined using high-performance liquid chromatography with tandem mass spectrometric detection (LC-MS/MS). Test Agent was isolated by liquid-liquid extraction. A partial aliquot of the supernatant was transferred to a clean 96-well collection plate, evaporated to dryness under nitrogen, and reconstituted with water. The extracted samples were analyzed using a Sciex 5500 mass spectrometer. The quantitative range of the assay was from 1-2,000 ng/mL.

Analysis of pharmacokinetic parameters – pharmacokinetic parameters were calculated using PkSolver^98^. Graphs were generated using PkSolver.

#### Additional Methods

(1) Enzyme inhibition assay, (2) Fluorescence/luminescence interference test, and (3) Colloidal aggregation study can be found in the Supporting Information.

## ASSOCIATED CONTENT

### Supporting Information

Molecular formula strings; Purification and characterization of all newly synthesized and purchased compounds (S1-S24); *in vitro* ADME (S25-S34); Additional Methods: Enzyme assay S35-S36); Colloidal aggregation study (S36-S37); Table S1 and S2 (S38); Figure S1 (S39); Figure S2 (S40); Figure S3 (S41).

## ACKNOWLEDGMENT

The study was supported by an intramural fund to Asim K Debnath (AKD) from the New York Blood Center. We gratefully acknowledge the generous gift of cells (highly expressed ACE2) and several plasmids to create the pseudovirus from Dr. Paul Bieniasz of Rockefeller University.

## ABBREVIATIONS USED

ACE2: Angiotensin-Converting Enzyme 2
ADME: Absorption, Distribution, Metabolism and Excretion
COVID-19: coronavirus disease 2019
CPE: Cytopathic Effect
HR1: Heptad Repeat 1
HR2: Heptad Repeat 2
MERS-CoV: Middle East Respiratory Syndrome-Corona Virus
RBD: Receptor Binding Domain
PFU: Plaque-Forming Unit
SARS-CoV: Severe Acute Respiratory Syndrome-Corona Virus.

## Table of Contents Graphics

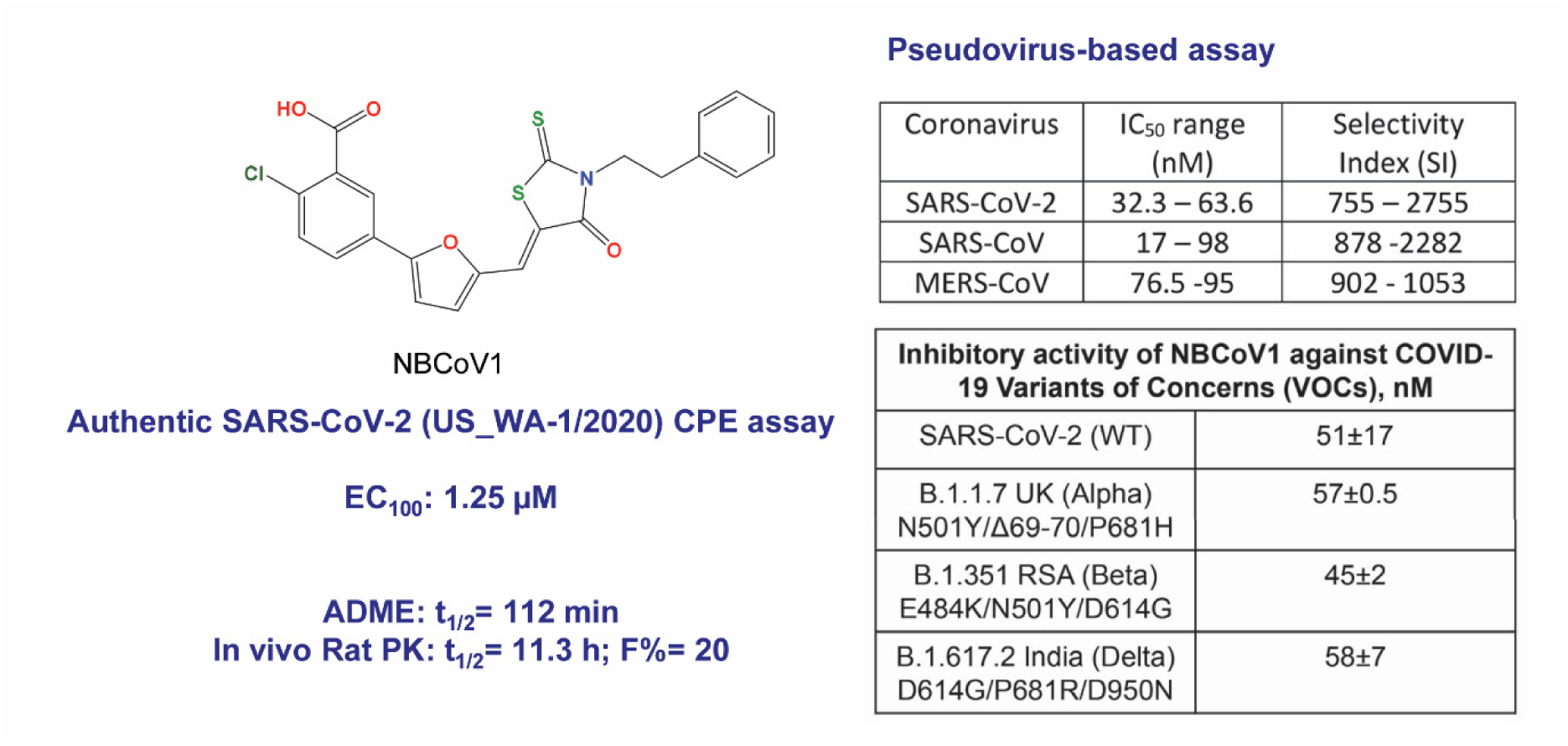

Pseudovirus-based assay
| Coronavirus | IC<sub>50</sub> range (nM) | Selectivity Index (SI) |
| --- | --- | --- |
| SARS-CoV-2 | 32.3 – 63.6 | 755 – 2755 |
| SARS-CoV | 17 – 98 | 878 -2282 |
| MERS-CoV | 76.5 -95 | 902 - 1053 |

Inhibitory activity of NBCoV1 against COVID-19 Variants of Concerns (VOCs), nM
|  |  |
| --- | --- |
| SARS-CoV-2 (WT) | 51±17 |
| B.1.1.7 UK (Alpha)<br>N501Y/Δ69-70/P681H | 57±0.5 |
| B.1.351 RSA (Beta)<br>E484K/N501Y/D614G | 45±2 |
| B.1.617.2 India (Delta)<br>D614G/P681R/D950N | 58±7 |

