## Supplementary material for "Discovery of highly potent pancoronavirus fusion inhibitors that also effectively inhibit COVID-19 variants from the UK (Alpha), South Africa (Beta), and India (Delta)": Table-S1

[*]Corresponding authors

Severe Acute Respiratory Syndrome (SARS), SARS-Cov-1, SARS-CoV-2, Middle East Respiratory Syndrome (MERS), MERS-CoV, COVID-19, pancoronavirus, fusion inhibitor

**List of Items**

**Purification and characterization of NBC0V1……………………………………………S3-S9**

**Analytical data on NBC0V15……………………………………………………………..S10-S12**

**Analytical data on NBCoV17……………………………………………………………..S13-S18**

**Analytical data on NBCoV28……………………………………………………………..S19-S21**

**Analytical data on NBCoV34……………………………………………………………..S22-S24**

**In vitro ADME………………………………………………………………………………S25-S34**

**Additional Methods……………………………………………………………………..S35-S37**

**Table-S1 and S2…………………………………………………………………………S38**

**FigureS1………………………………………………………………………………………..S39**

**FigureS2………………………………………………………………………………………..S40**

**FigureS3………………………………………………………………………………………..S41**

#### Experimental Section

##### Purification and characterization of (Z)-2-Chloro-5-(5-((4-oxo-3-phenethyl-2-thioxothiazolidin-5-ylidene)methyl)furan-2-yl)benzoic acid (NBCoV1) purchased from Sigma-Aldrich.

Common reagents and solvents were purchased from commercial suppliers and used without further purification unless otherwise stated. Tetrahydrofuran (THF) was distilled from sodium-benzophenone under an argon atmosphere. Reaction progress was monitored using analytical thin-layer chromatography (TLC) on pre-coated silica gel GF254 plates (Macherey Nagel GmbH & Co. KG, Düren, Germany), and spots were detected under UV light (254 and 366 nm). Compounds were purified with flash column chromatography with a silica gel and particle size of 40–63 μM (Merck, Darmstadt, Germany) as the stationary phase and hexane/ethyl acetate or dichloromethane/methanol mixtures as eluent systems. Nuclear magnetic resonance spectra were measured on a Bruker AV-400 NMR instrument (Bruker, Karlsruhe, Germany) in deuterated solvents (DMSO-$d_6$, $CDCl_3$, MeOD-$d_4$). Chemical shifts are expressed in ppm relative to DMSO-$d_6$ or MeOD-$d_4$ (2.50/3.31 for 1H; 39.52/49.00 for 13C). The following abbreviations are used to set multiplicities: s = singlet, d = doublet, t = triplet, q = quartet, m = multiplet, br. = broad. Measurements for verification and purity of the compounds were performed by LC/MS. LC–MS/MS data were obtained using a Dionex Ultimate 3000 liquid chromatograph (Dionex, USA) connected to an AB Sciex Qtrap 3200 mass spectrometer (AB Sciex, Canada). LC separation was carried out on a Shim-pack GIST C18-AQ (150 mm × 2.1 mm, 3 μm, Shimadzu, Japan) column. Mobile phase consisted of the mixture of 0.1% (v/v) formic acid in water (A) and acetonitrile (B). Separation was performed in isocratic mode 10% : 90% (A : B). The mobile phase flow rate was 0.3 mL $min^{-1}$. The injection volume was 10 μL. Compounds were detected at λ =

254 nm. All high resolution mass spectra (HRMS) were measured on AB Sciex TripleTOF 5600+ instrument equipped with DuoSpray (ESI) ion source. Samples were directly injected in the ion source in acetonitrile or methanol solutions acidified by formic acid. The melting points were measured in open capillaries and presented without correction.

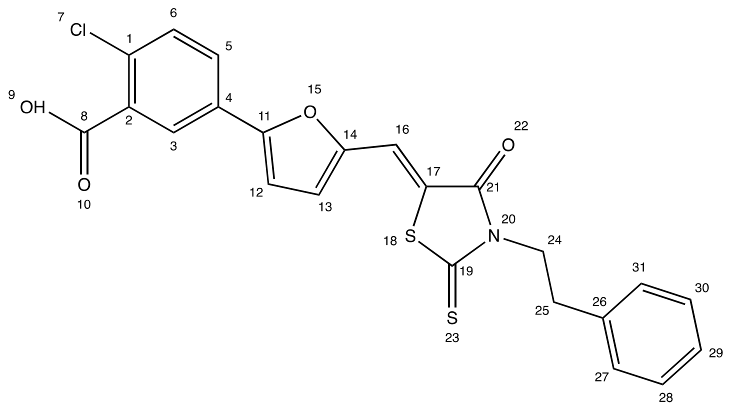

**(Z)-2-Chloro-5-(5-((4-oxo-3-phenethyl-2-thioxothiazolidin-5-ylidene)methyl)furan-2-yl)benzoic acid**

rt = 2.087 min. Purity = 98%. Mp = 348 – 350 °C. LC–MS: m/z [M+Na]+ = 492 Da.

**$^1$H NMR (400.13 MHz, DMSO-d$_6$): δ** = 7.79 (d, *J* = 2.4 Hz, 1 H, 6), 7.64 (s, 1 H, 3), 7.62 (dd, *J* = 8.4 Hz, 2.2 Hz, 1 H, 5), 7.42 – 7.47 (m, *J* = 3.5 Hz, 1 H, 16), 7.37 (d, *J* = 3.5 Hz, 1 H, 13), 7.34 (d, *J* = 3.5 Hz, 1 H, 12), 7.25 – 7.35 (m, 2 H, 27, 31), 7.18 – 7.24 (m, 3 H, 28, 29, 30), 4.23 - 4.32 (m, 2 H, 24), 2.90 - 3.02 (m, 2 H, 25).

**$^{13}$C NMR (100.61 MHz, DMSO-*d*$_6$)**: **δ =** 193.88 (s, 1 C, 19), 168.94 (s, 1 C, 8), 166.56 (s, 1 C, 21), 157.82 (s, 1 C, 11), 149.46 (s, 1 C, 14), 144.65 (s, 1 C, 2), 137.87 (s, 1 C, 1), 130.41 (s, 1 C, 3), 130.13 (s, 1 C, 26), 128.93 (s, 2 C, 28, 30), 128.77 (s, 2 C, 27, 31), 126.85 (s, 1 C, 29), 124.31 (s, 1 C, 6), 123.66 (s, 1 C, 5), 123.46 (s, 1 C, 17), 123.27 (s, 1 C, 4), 118.44 (s, 1 C, 13), 118.43 (s, 1 C, 12), 110.71 (s, 1 C, 16), 45.50 (s, 1 C, 24), 32.37 (s, 1 C, 25).

**HRMS (ESI):** for $C_{23}H_{16}ClNO_4S_2$ $[M+Na]^+$: calcd 492.0102, found 492.0106

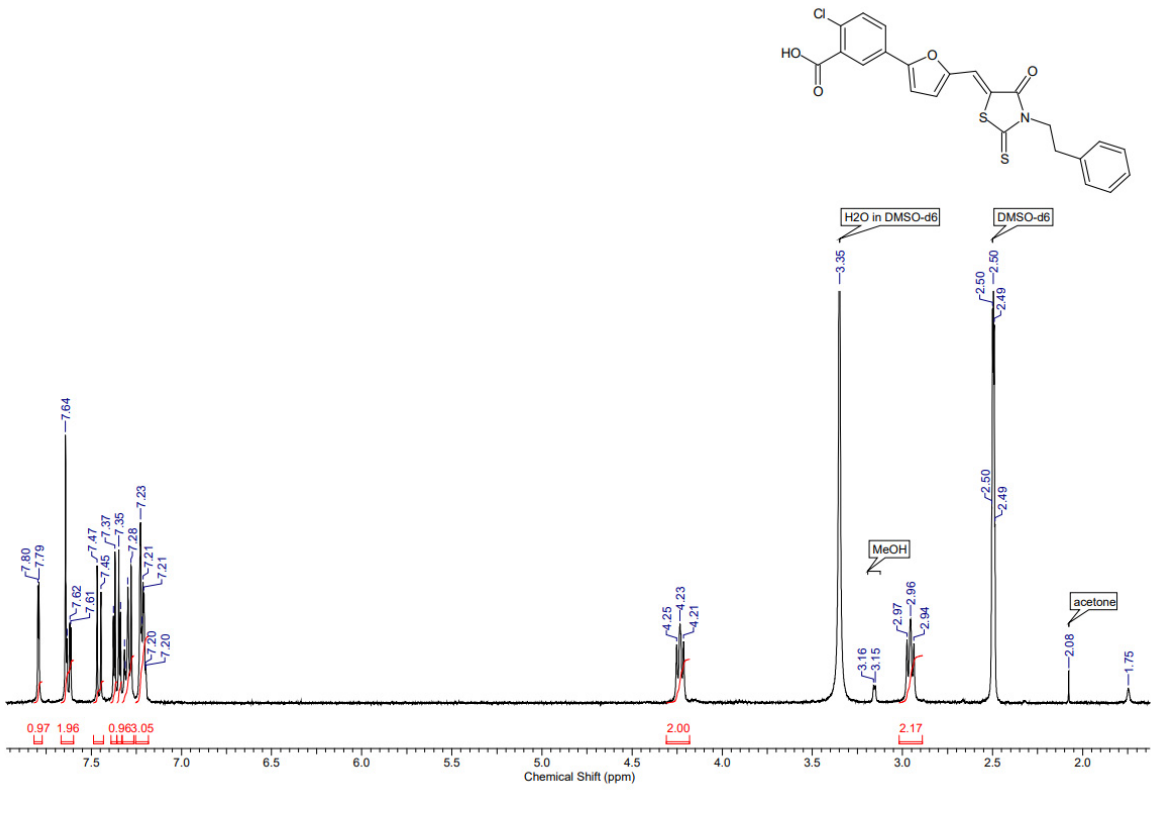

Cl
HO
O
O
O
S
N
S
H2O in DMSO-d6
DMSO-d6
MeOH
acetone
7.80
7.79
7.64
7.62
7.61
7.47
7.45
7.37
7.35
7.28
7.23
7.21
7.21
7.20
7.20
4.25
4.23
4.21
3.35
3.16
3.15
2.97
2.96
2.94
2.50
2.50
2.49
2.50
2.49
2.08
1.75
0.97
1.96
0.96
3.05
2.00
2.17
7.5
7.0
6.5
6.0
5.5
5.0
4.5
4.0
3.5
3.0
2.5
2.0
Chemical Shift (ppm)

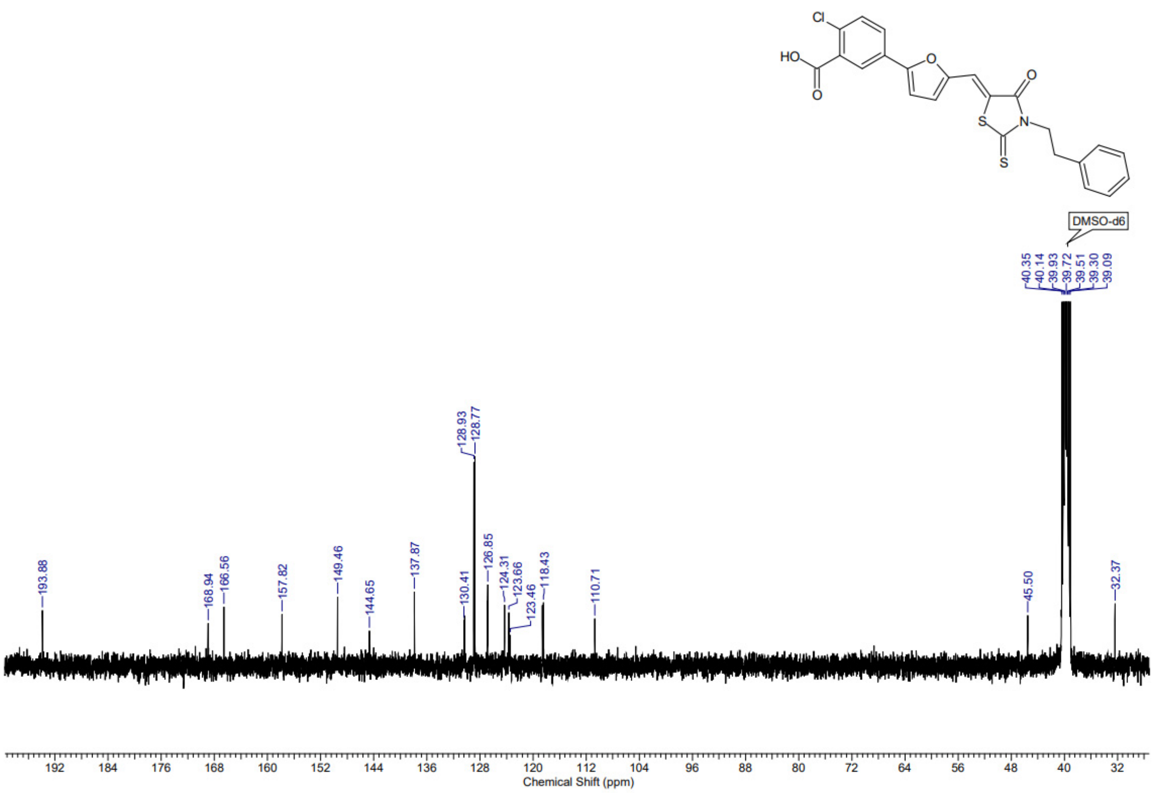
Cl
HO
O
O
S
N
O
S
DMSO-d6
40.35
40.14
39.93
39.72
39.51
39.30
39.09
128.93
128.77
193.88
168.94
166.56
157.82
149.46
144.65
137.87
130.41
126.85
124.31
123.66
123.46
118.43
110.71
45.50
32.37
192 184 176 168 160 152 144 136 128 120 112 104 96 88 80 72 64 56 48 40 32
Chemical Shift (ppm)

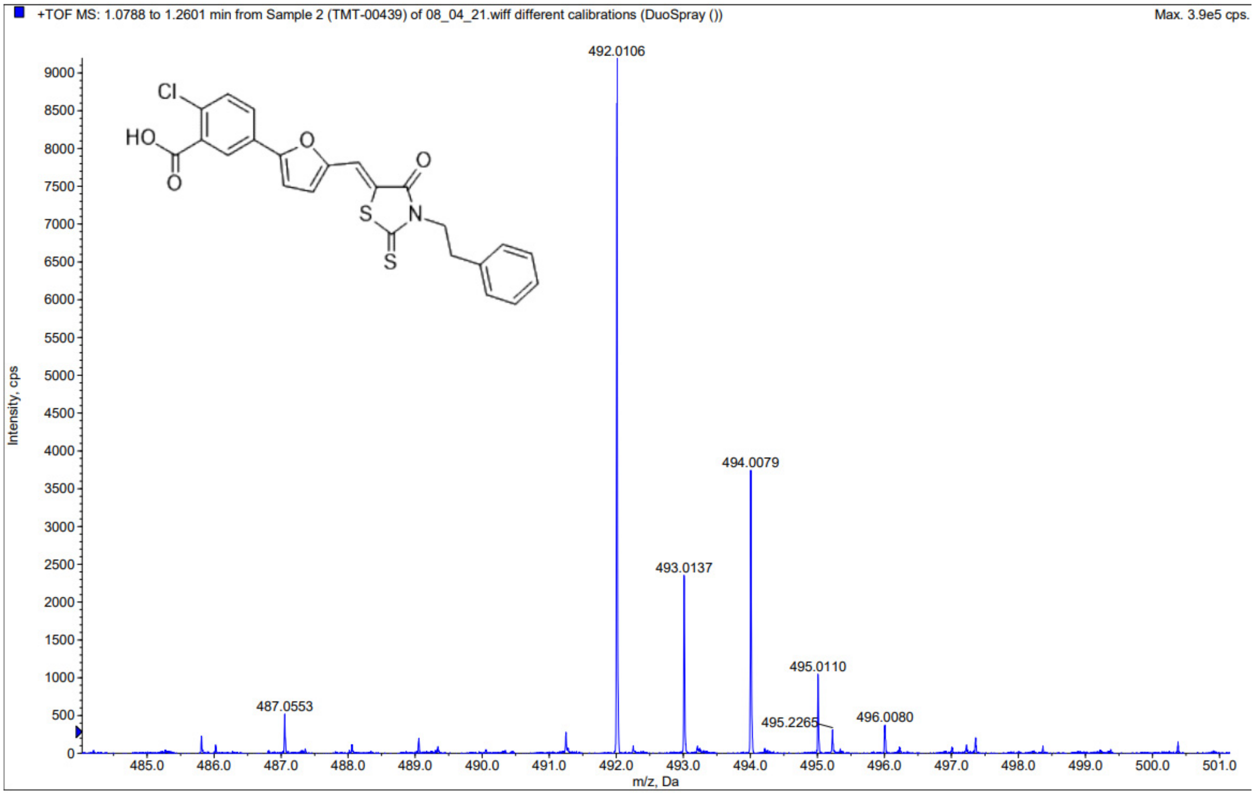
+TOF MS: 1.0788 to 1.2601 min from Sample 2 (TMT-00439) of 08_04_21.wiff different calibrations (DuoSpray ())
Max. 3.9e5 cps.
492.0106
494.0079
493.0137
495.0110
495.2265
496.0080
487.0553
Intensity, cps
m/z, Da
Cl
HO
O
O
O
S
N
S

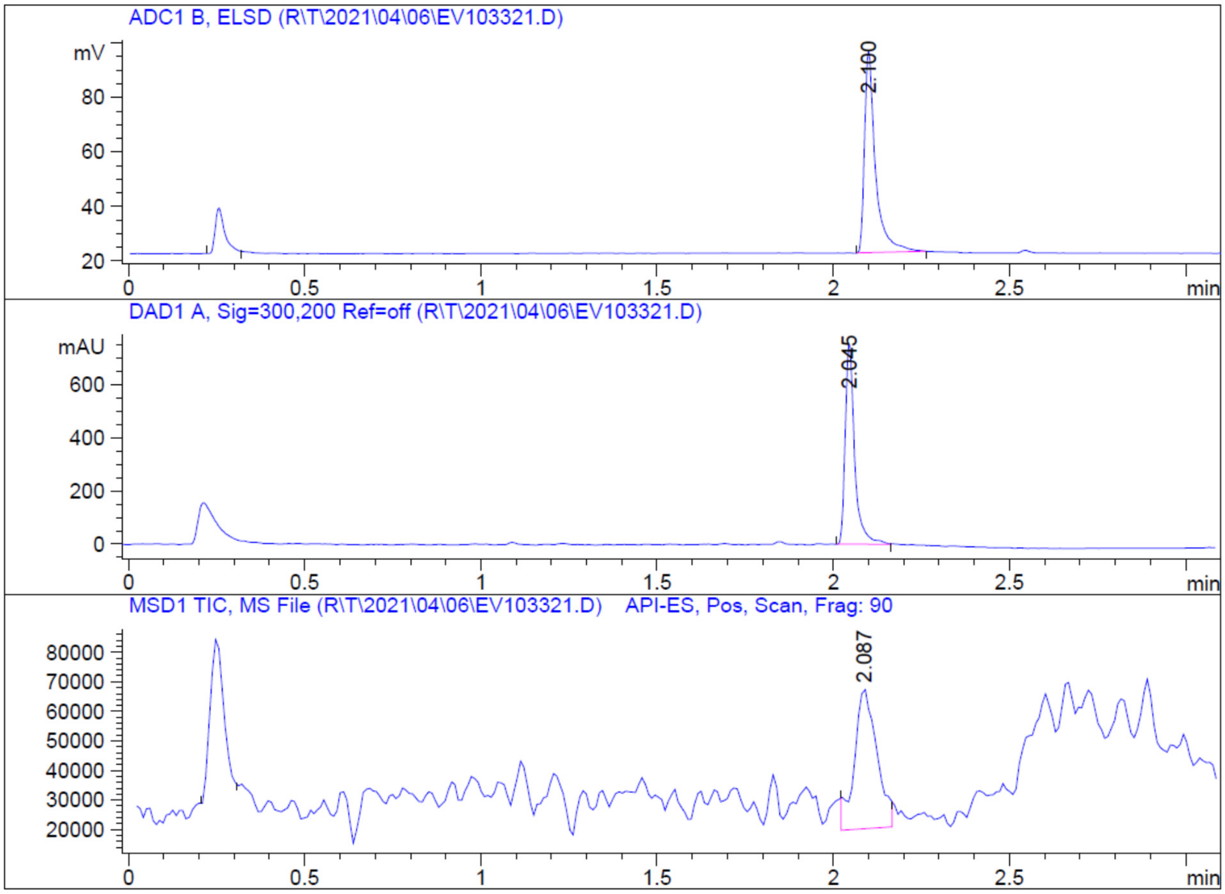
ADC1 B, ELSD (R\T\2021\04\06\EV103321.D)
mV
2.100
DAD1 A, Sig=300,200 Ref=off (R\T\2021\04\06\EV103321.D)
mAU
2.045
MSD1 TIC, MS File (R\T\2021\04\06\EV103321.D) API-ES, Pos, Scan, Frag: 90
2.087
min

**Analytical data on NBCoV15 purchased from Chembridge Corp (San Diego, CA)**

Purity: 100%. MW: 425.9593 (Calc.); 425.9 (found).

CR00902482

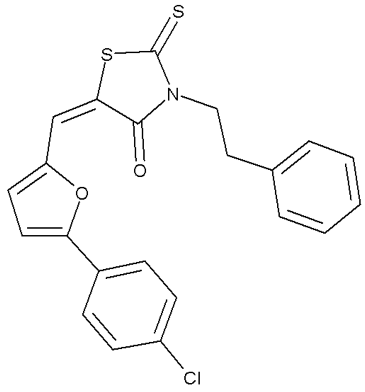

| | | | |
|---|---|---|---|
| ID | 5521729 | 425.9593 | $C_{22}H_{16}ClNO_2S_2$ |

Data File R:\HPLC\AUTO\CR009024\1BK-0901.D
Sample Name: CR009024P1-B-11
Instrument 1 15/04/2021 12:48:19 1
Column: ONYX MONOLITHIC, C18 50x3mm |1.8ml/min| Columns Reg Valve
Gradient: "A"->@2.0min->"B"(Hold 0.6min)->@0.2min->"A"->PostRun
PMP1, Solvent A : 0.1%TFA, 2.5%AcN in H2O
PMP1, Solvent B : 0.1%TFA in AcN
PMP1, Solvent C : --NOT USED--
PMP1, Solvent D : --NOT USED--
Ionization mode : API-ES Positive

Signal 1: ADC1 B, ELSD

```
Peak RetTime Type  Width     Area      Height     Area
  #   [min]        [min]   [mV*s]      [mV]         %
----|-------|----|-------|----------|----------|--------|
   1   2.458 PPA   0.0312    3.15085    1.51854 100.0000
Totals :                     3.15085    1.51854
```

Signal 2: DAD1 A, Sig=300,200 Ref=off

```
Peak RetTime Type  Width     Area      Height     Area
  #   [min]        [min]   [mAU*s]     [mAU]        %
----|-------|----|-------|----------|----------|--------|
   1   2.404 BP    0.0299  113.99640   58.00123 100.0000
Totals :                   113.99640   58.00123
```

Signal 3: MSD1 TIC, MS File

| Peak # | RetTime [min] | Type | Width [min] | Area | Height | Area % |
|---|---|---|---|---|---|---|
| 1 | 2.448 | MM | 0.0526 | 8.40224e4 | 2.66246e4 | 100.0000 |
| Totals : | | | | 8.40224e4 | 2.66246e4 | |

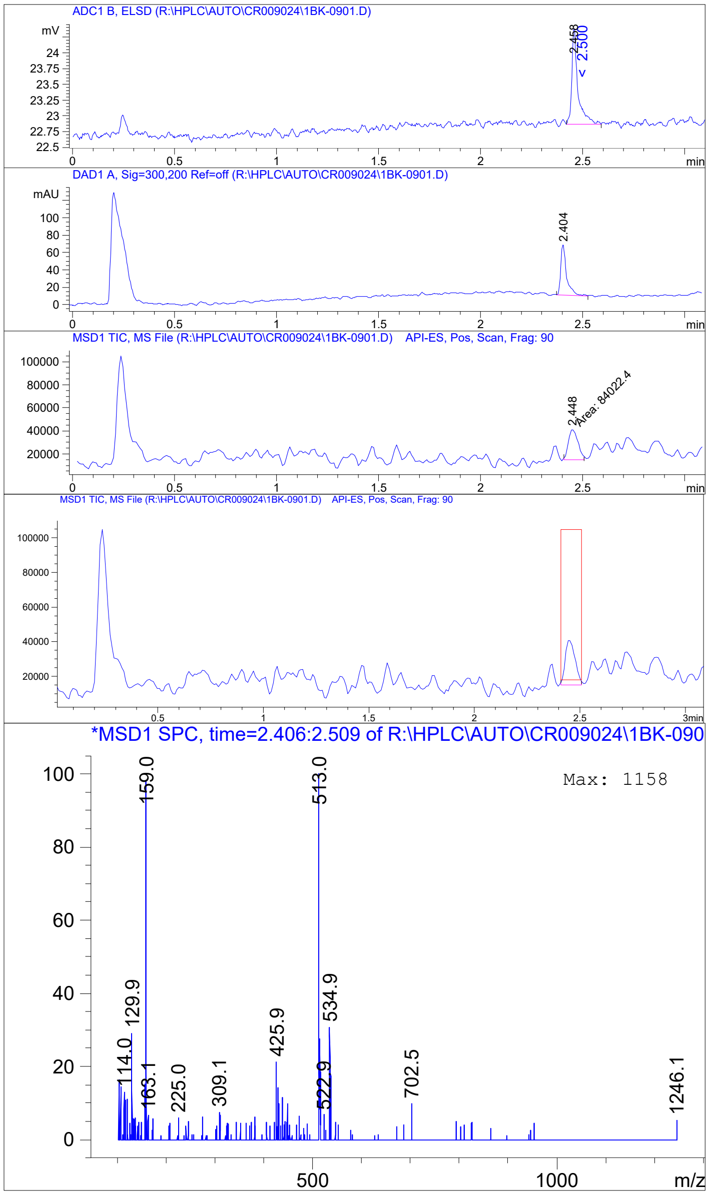

ADC1 B, ELSD (R:\HPLC\AUTO\CR009024\1BK-0901.D)
DAD1 A, Sig=300,200 Ref=off (R:\HPLC\AUTO\CR009024\1BK-0901.D)
MSD1 TIC, MS File (R:\HPLC\AUTO\CR009024\1BK-0901.D) API-ES, Pos, Scan, Frag: 90
Area: 84022.4
*MSD1 SPC, time=2.406:2.509 of R:\HPLC\AUTO\CR009024\1BK-090
Max: 1158
m/z

521729x2 in DMSO.

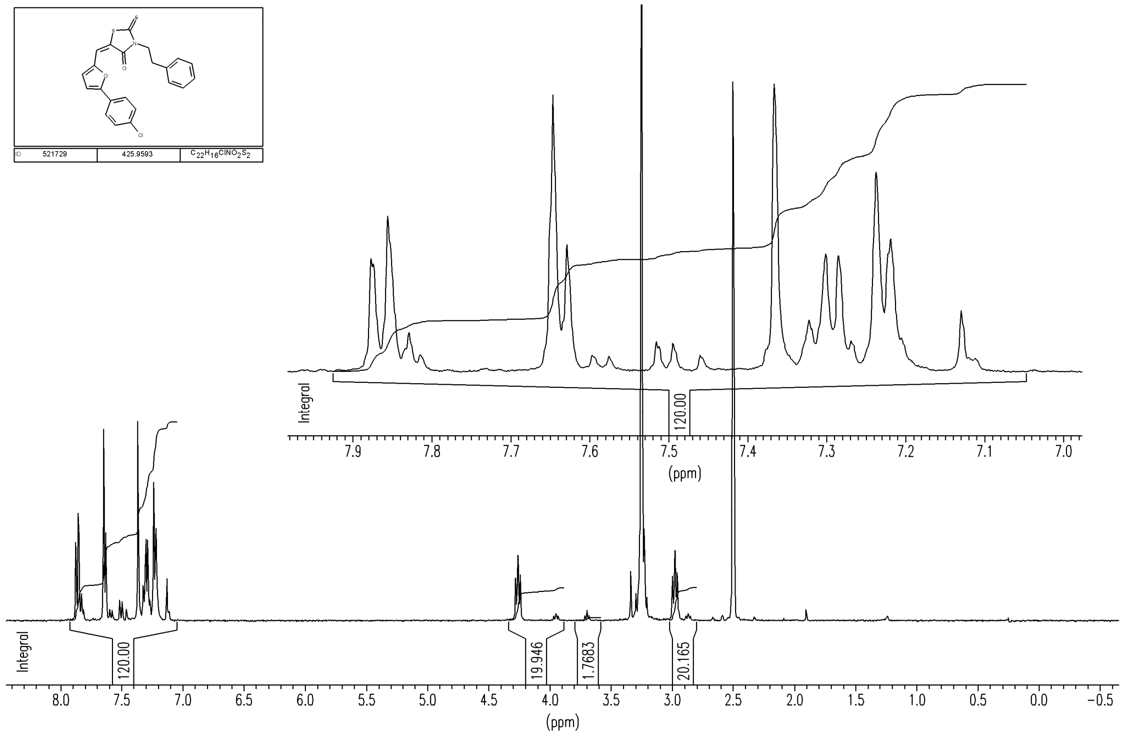

**Analytical data on NBCoV17 purchased from Chembridge Corp (San Diego, CA)**

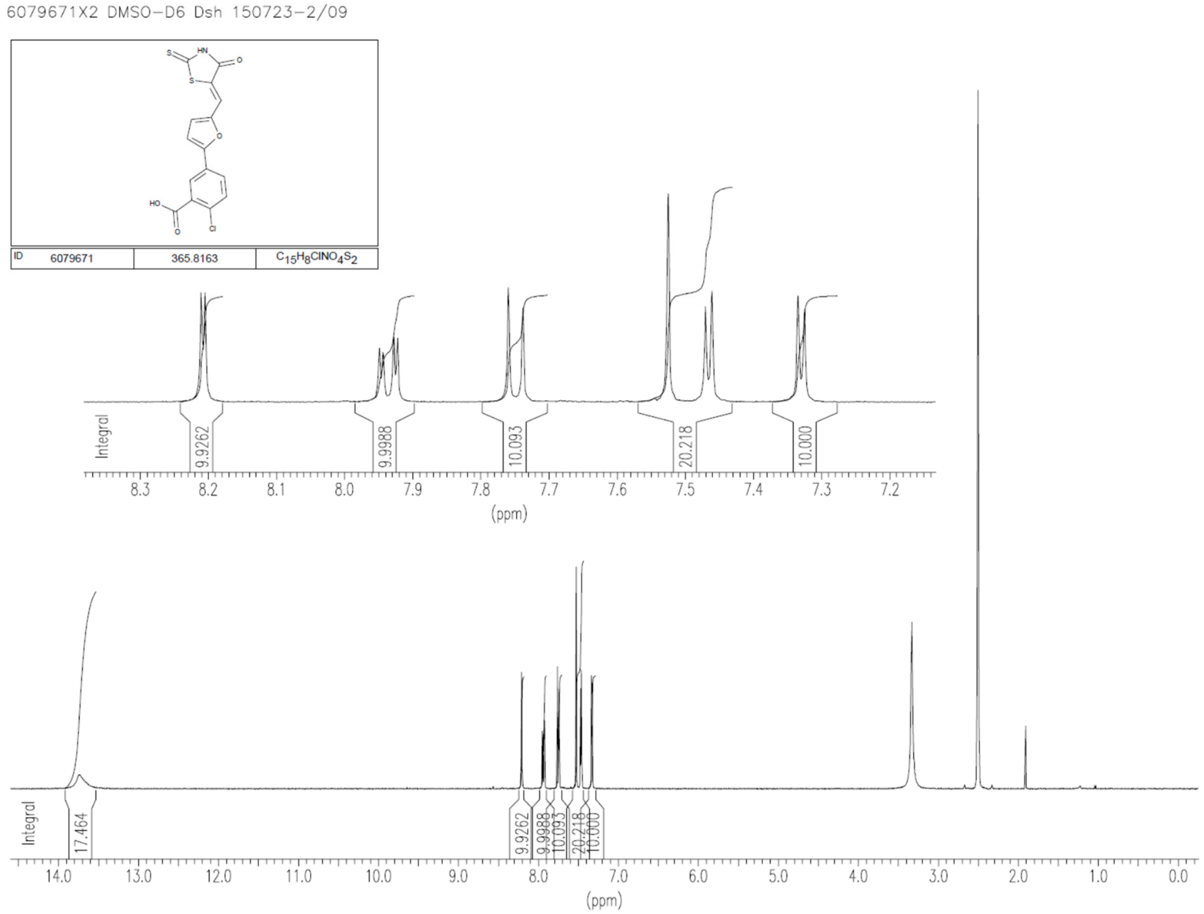

CR00900548

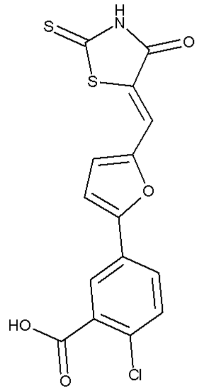

| | | |
|---|---|---|
| ID 6079671 | 365.8163 | $C_{15}H_8ClNO_4S_2$ |

Data File D:\DATA\MICRA\2HF-0101.D

Sample Name: CR009005P2-H-06
Instrument 1 23/07/2015 16:34:24 6
Column: Onyx Monolithic C18 50x4.6mm | 3.75ml/min | Columns Reg Valve
Gradient: "C"->@2.2min->"D"(Hold 0.4min)->@0.2min->"C"->PostRun
PMP1, Solvent A : 0.1%TFA in Acn/H2O (2.5:97.5)
PMP1, Solvent B : 0.1%TFA in AcN
PMP1, Solvent C : 0.1%FA in Acn/H2O (2.5:97.5)
PMP1, Solvent D : 0.1%FA in AcN
Ionization mode : APCI Negative

Signal 1: ADC1 B, ELSD

| Peak # | RetTime [min] | Type | Width [min] | Area [mAu*s] | Height [mAu] | Area % |
|---|---|---|---|---|---|---|
| 1 | 5.556 | MM | 0.0550 | 22.90727 | 6.93644 | 100.0000 |
| Totals : | | | | 22.90727 | 6.93644 | |

Signal 2: DAD1 A, Sig=300,200 Ref=off

| Peak # | RetTime [min] | Type | Width [min] | Area [mAU*s] | Height [mAU] | Area % |
|---|---|---|---|---|---|---|
| 1 | 0.709 | MM | 0.0889 | 47.11327 | 8.83276 | 4.0217 |
| 2 | 5.512 | MM | 0.0626 | 1124.37195 | 299.48669 | 95.9783 |
| Totals : | | | | 1171.48522 | 308.31946 | |

Signal 3: MSD1 TIC, MS File

| Peak # | RetTime [min] | Type | Width [min] | Area | Height | Area % |
|---|---|---|---|---|---|---|
| 1 | 5.539 | MM | 0.0744 | 2.62775e6 | 5.89026e5 | 100.0000 |
| Totals : | | | | 2.62775e6 | 5.89026e5 | |

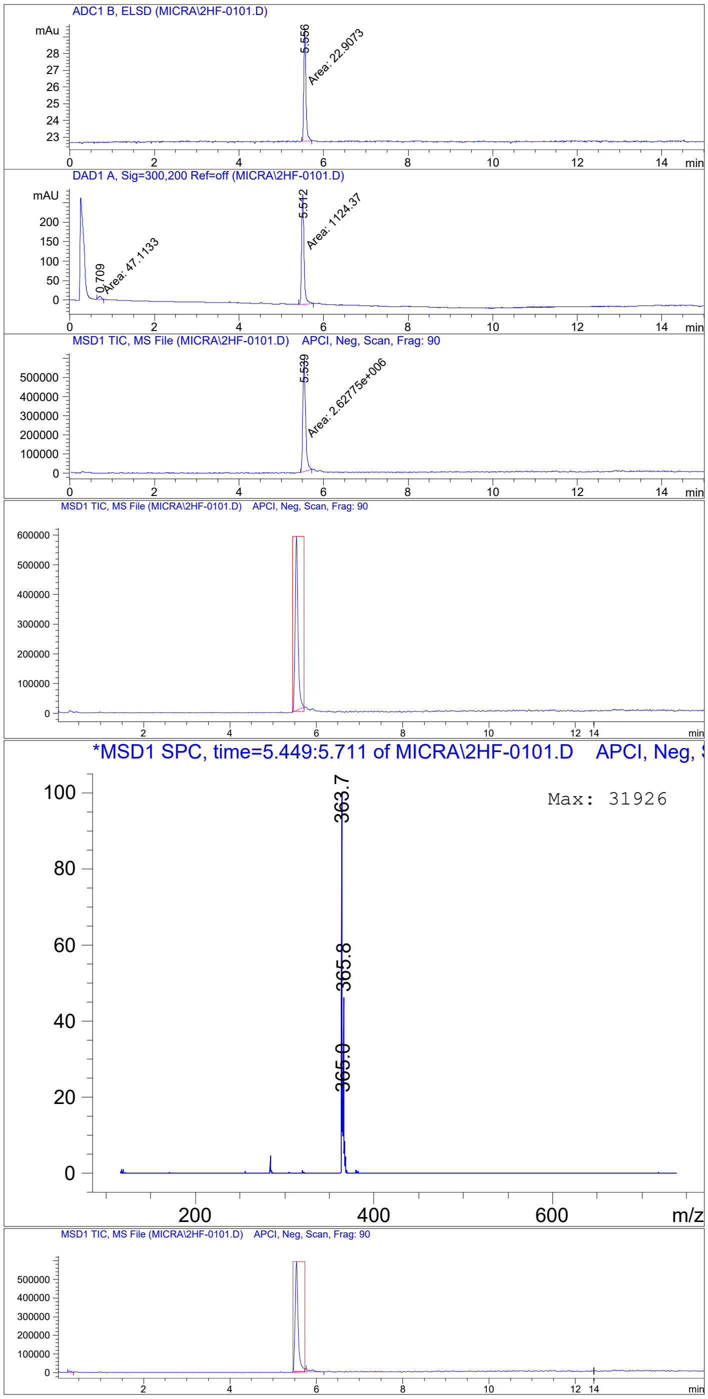
ADC1 B, ELSD (MICRA\2HF-0101.D)
5.556
Area: 22.9073
DAD1 A, Sig=300,200 Ref=off (MICRA\2HF-0101.D)
0.709
Area: 47.1133
5.512
Area: 1124.37
MSD1 TIC, MS File (MICRA\2HF-0101.D) APCI, Neg, Scan, Frag: 90
5.539
Area: 2.62775e+006
MSD1 TIC, MS File (MICRA\2HF-0101.D) APCI, Neg, Scan, Frag: 90
*MSD1 SPC, time=5.449:5.711 of MICRA\2HF-0101.D APCI, Neg,
Max: 31926
363.7
365.8
365.0
m/z
MSD1 TIC, MS File (MICRA\2HF-0101.D) APCI, Neg, Scan, Frag: 90

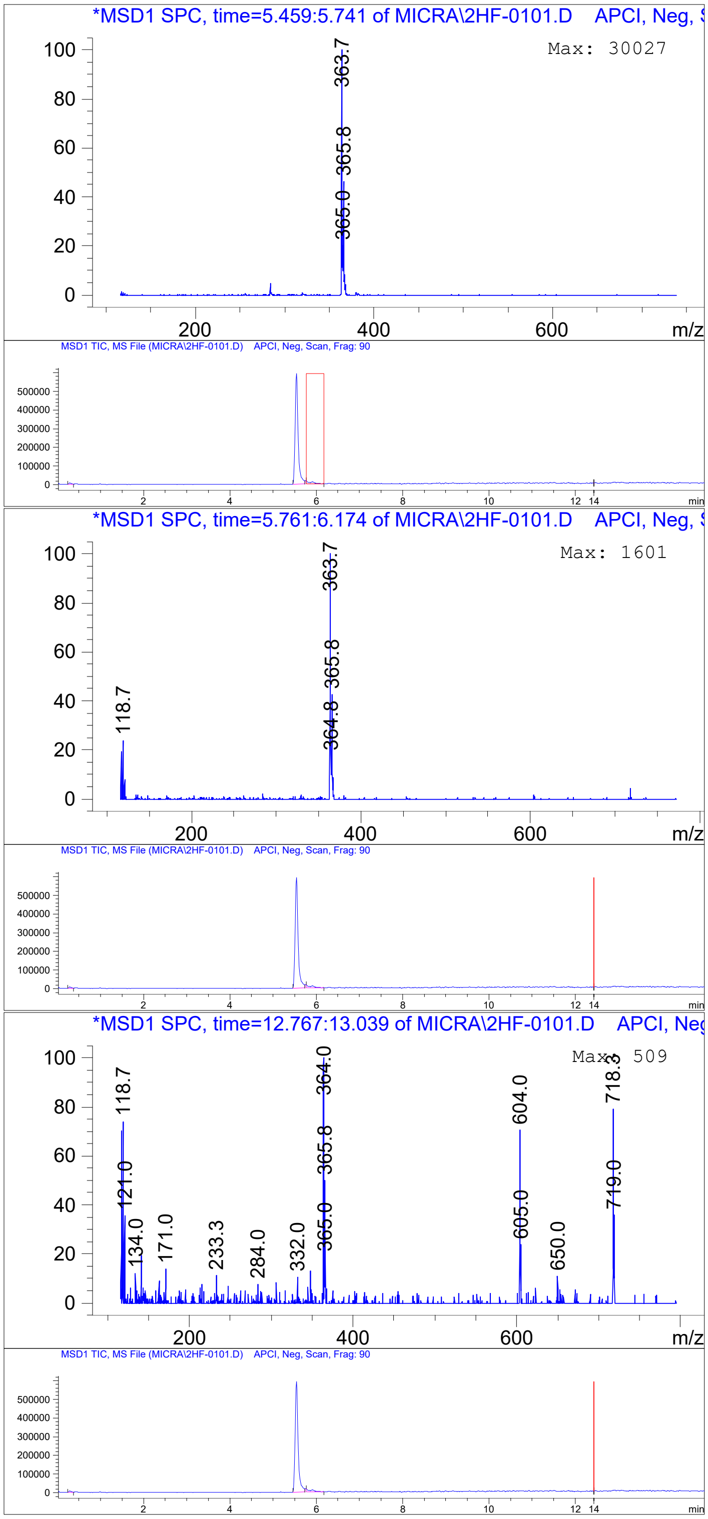
*MSD1 SPC, time=5.459:5.741 of MICRA\2HF-0101.D APCI, Neg,
Max: 30027
363.7
365.8
365.0
m/z
MSD1 TIC, MS File (MICRA\2HF-0101.D) APCI, Neg, Scan, Frag: 90
min
*MSD1 SPC, time=5.761:6.174 of MICRA\2HF-0101.D APCI, Neg,
Max: 1601
118.7
363.7
365.8
364.8
m/z
MSD1 TIC, MS File (MICRA\2HF-0101.D) APCI, Neg, Scan, Frag: 90
min
*MSD1 SPC, time=12.767:13.039 of MICRA\2HF-0101.D APCI, Neg
Max: 509
118.7
121.0
134.0
171.0
233.3
284.0
332.0
364.0
365.8
365.0
604.0
605.0
650.0
718.3
719.0
m/z
MSD1 TIC, MS File (MICRA\2HF-0101.D) APCI, Neg, Scan, Frag: 90
min

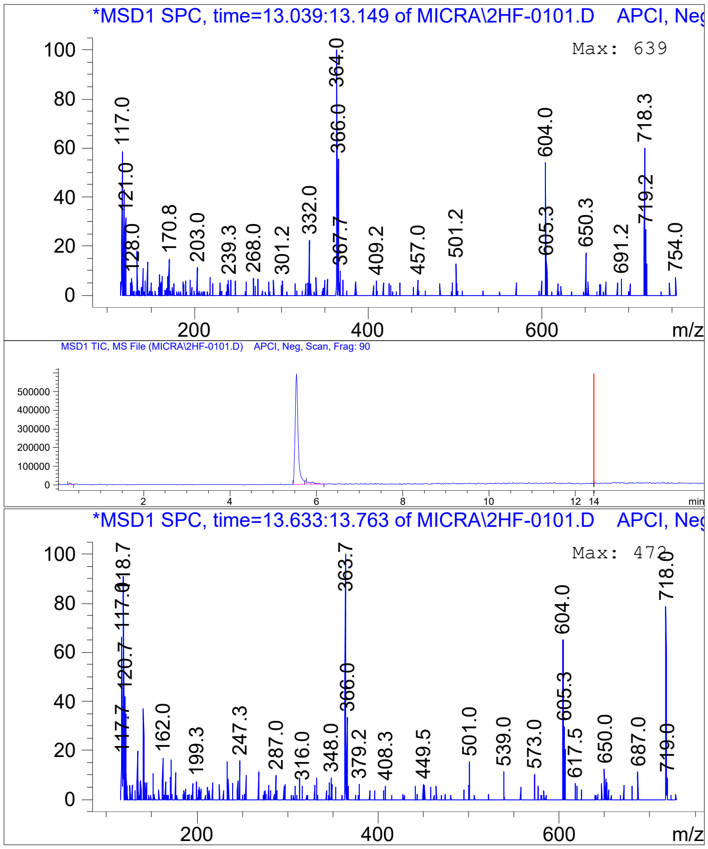
*MSD1 SPC, time=13.039:13.149 of MICRA\2HF-0101.D APCI, Neg
Max: 639
MSD1 TIC, MS File (MICRA\2HF-0101.D) APCI, Neg, Scan, Frag: 90
*MSD1 SPC, time=13.633:13.763 of MICRA\2HF-0101.D APCI, Neg
Max: 472
m/z
min

**Analytical data on NBCoV28 purchased from Chembridge Corp (San Diego, CA)**

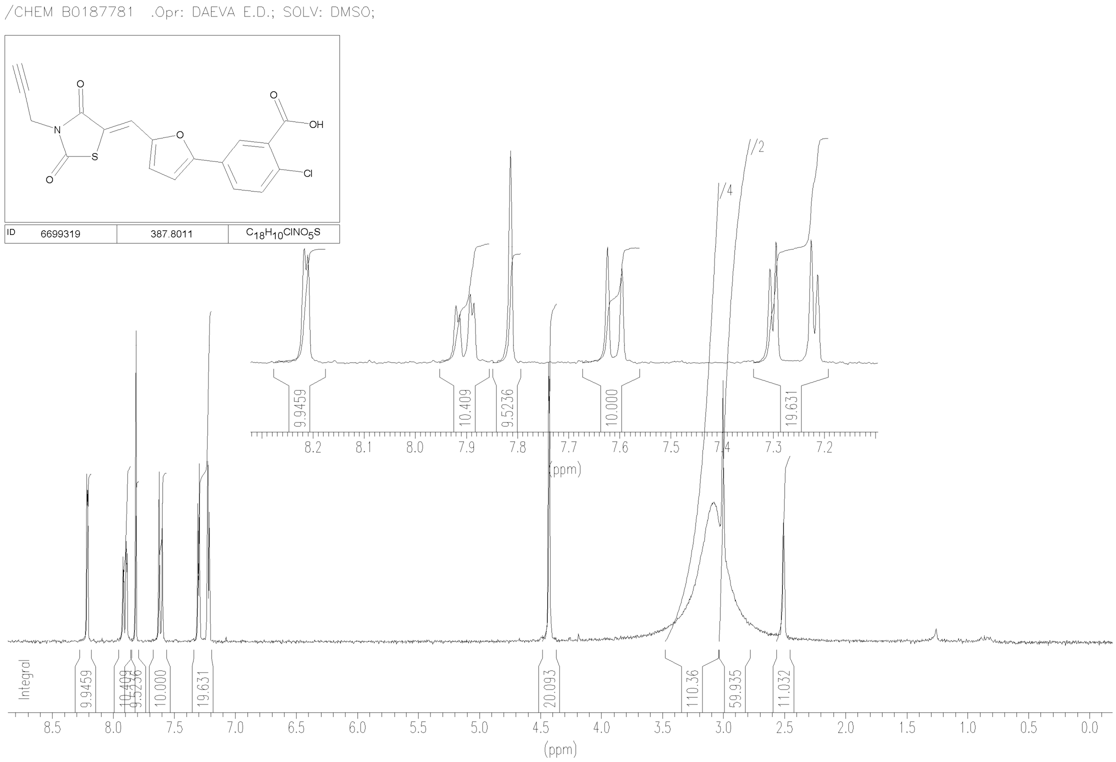

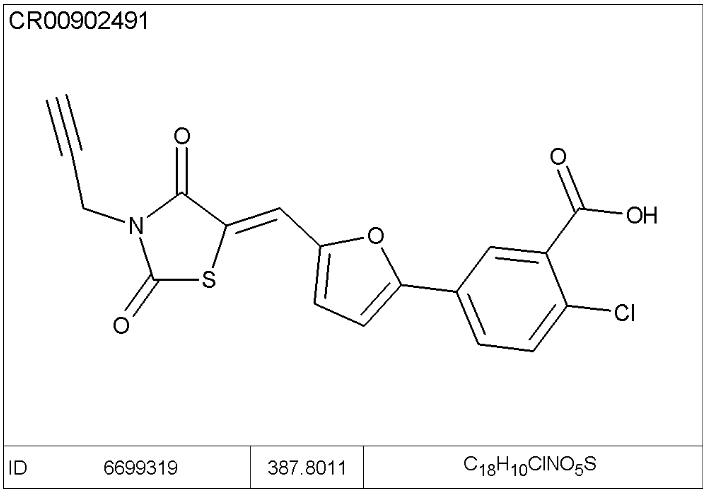

Data File R:\HPLC\AUTO\CR009024\1CL-0901.D

Sample Name: CR009024P1-C-12
Instrument 1 20/04/2021 14:50:59 1
Column: Onyx C18 50x4.6mm | 3.75ml/min | Columns Reg Valve
Gradient: "A"->@2.2min->"B"(Hold 0.4min)->@0.2min->"A"->PostRun
PMP1, Solvent A : 0.1TFA in 2.5%AcN/W
PMP1, Solvent B : 0.1TFA in AcN
PMP1, Solvent C : 0.1FA in 2.5%AcN/W
PMP1, Solvent D : 0.1FA in AcN
Ionization mode : API-ES Negative

Signal 1: ADC1 A, ADC1 ELSD

| Peak # | RetTime [min] | Type | Width [min] | Area [mAU*s] | Height [mAU] | Area % |
|---|---|---|---|---|---|---|
| 1 | 1.572 | MM | 0.0317 | 28.71860 | 15.10031 | 92.2487 |
| 2 | 2.421 | MM | 0.0276 | 2.41310 | 1.45476 | 7.7513 |
| Totals : | | | | 31.13171 | 16.55507 | |

Signal 2: DAD1 A, Sig=300,200 Ref=off

| Peak # | RetTime [min] | Type | Width [min] | Area [mAU*s] | Height [mAU] | Area % |
|---|---|---|---|---|---|---|
| 1 | 1.523 | MF | 0.0248 | 575.02313 | 386.74631 | 93.7920 |
| 2 | 1.554 | FM | 0.0211 | 38.06044 | 30.04487 | 6.2080 |
| Totals : | | | | 613.08357 | 416.79118 | |

Signal 3: MSD1 TIC, MS File

| Peak # | RetTime [min] | Type | Width [min] | Area | Height | Area % |
|---|---|---|---|---|---|---|
| 1 | 1.536 | BP | 0.0365 | 2.58954e6 | 1.10936e6 | 100.0000 |
| Totals : | | | | 2.58954e6 | 1.10936e6 | |

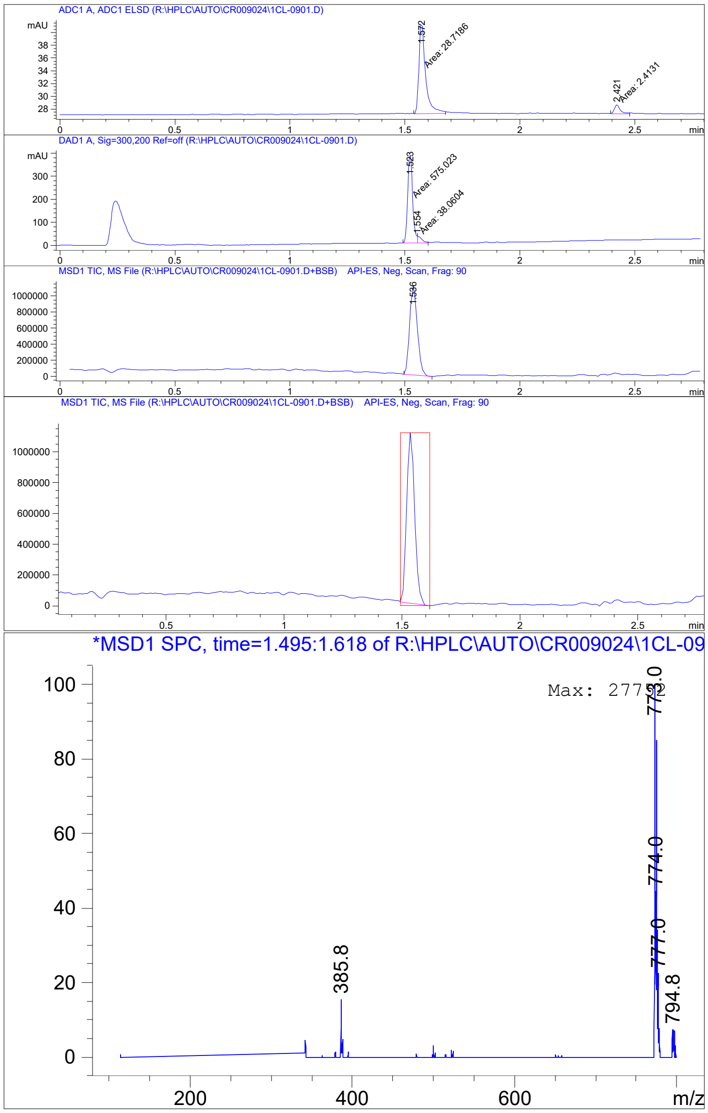
ADC1 A, ADC1 ELSD (R:\HPLC\AUTO\CR009024\1CL-0901.D)
Area: 28.7186
Area: 2.4131
DAD1 A, Sig=300,200 Ref=off (R:\HPLC\AUTO\CR009024\1CL-0901.D)
Area: 575.023
Area: 38.0604
MSD1 TIC, MS File (R:\HPLC\AUTO\CR009024\1CL-0901.D+BSB) API-ES, Neg, Scan, Frag: 90
MSD1 TIC, MS File (R:\HPLC\AUTO\CR009024\1CL-0901.D+BSB) API-ES, Neg, Scan, Frag: 90
*MSD1 SPC, time=1.495:1.618 of R:\HPLC\AUTO\CR009024\1CL-09
385.8
774.0
777.0
794.8
m/z

#### Analytical data on NBCoV34 purchased from Chembridge Corp (San Diego, CA)

660015-A

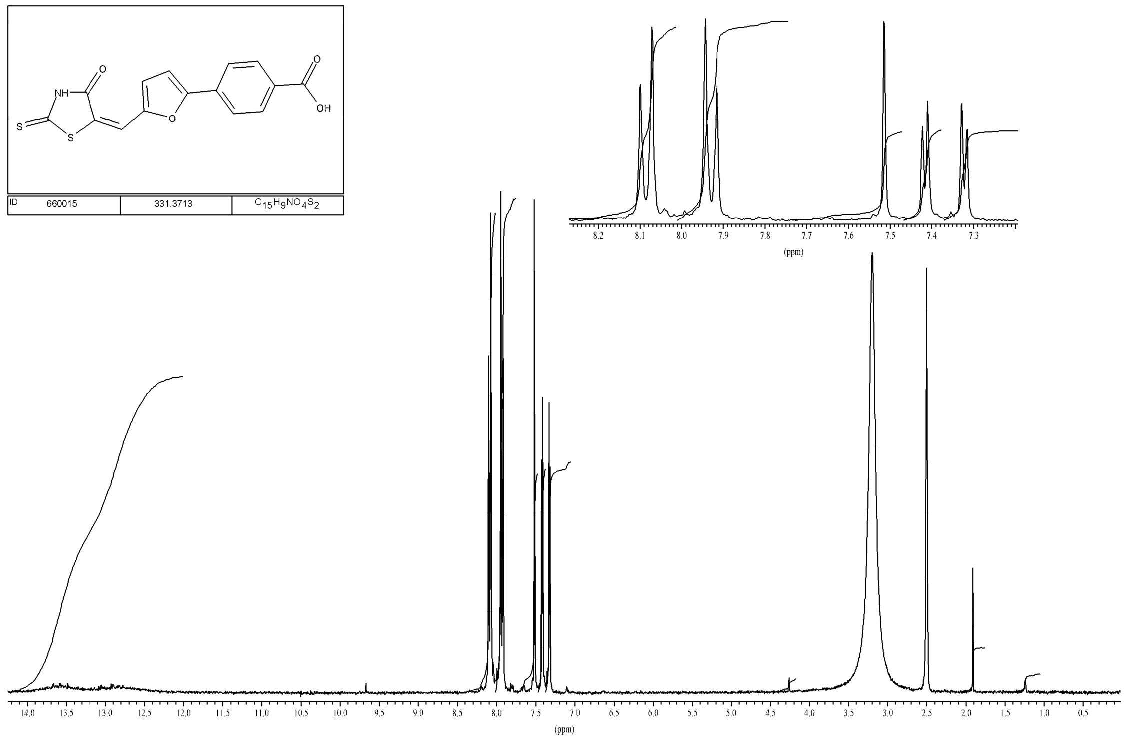

CR00902485

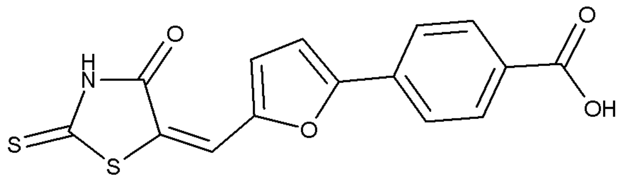

| ID | 5660015 | 331.3713 | $C_{15}H_9NO_4S_2$ |
|---|---|---|---|

Data File D:\DATA\MICRA\1EK-0301.D
Sample Name: CR009024P1-E-11
Instrument 1 20/04/2021 16:54:25 1
Column: Onyx C18 50x4.6mm | 3.75ml/min | Columns Reg Valve
Gradient: "A"->@2.0min->"B"(Hold 0.4min)->@0.2min->"A"->PostRun
PMP1, Solvent A : 0.1TFA in 2.5%AcN/W
PMP1, Solvent B : 0.1TFA in AcN
PMP1, Solvent C : 0.1FA in 2.5%AcN/W
PMP1, Solvent D : 0.1FA in AcN
Ionization mode : API-ES Negative

Signal 1: ADC1 A, ADC1 ELSD

| Peak # | RetTime [min] | Type | Width [min] | Area [mAU*s] | Height [mAU] | Area % |
|---|---|---|---|---|---|---|
| 1 | 1.463 | MM | 0.0391 | 1.73849 | 7.40851e-1 | 91.3606 |
| 2 | 2.409 | MM | 0.0206 | 1.64398e-1 | 1.32726e-1 | 8.6394 |
| Totals : | | | | 1.90288 | 8.73577e-1 | |

Signal 2: DAD1 A, Sig=300,200 Ref=off

| Peak # | RetTime [min] | Type | Width [min] | Area [mAU*s] | Height [mAU] | Area % |
|---|---|---|---|---|---|---|
| 1 | 1.425 | MF | 0.0366 | 150.02690 | 68.30225 | 93.9473 |
| 2 | 1.483 | FM | 0.0217 | 9.66572 | 7.43823 | 6.0527 |
| Totals : | | | | 159.69263 | 75.74048 | |

Signal 3: MSD1 TIC, MS File

| Peak # | RetTime [min] | Type | Width [min] | Area | Height | Area % |
|---|---|---|---|---|---|---|

| | | | | | | |
|---|---|---|---|---|---|---|
| 1 | 1.420 | MM | 0.0429 | 3.85948e5 | 1.49882e5 | 100.0000 |
| Totals : | | | | 3.85948e5 | 1.49882e5 | |

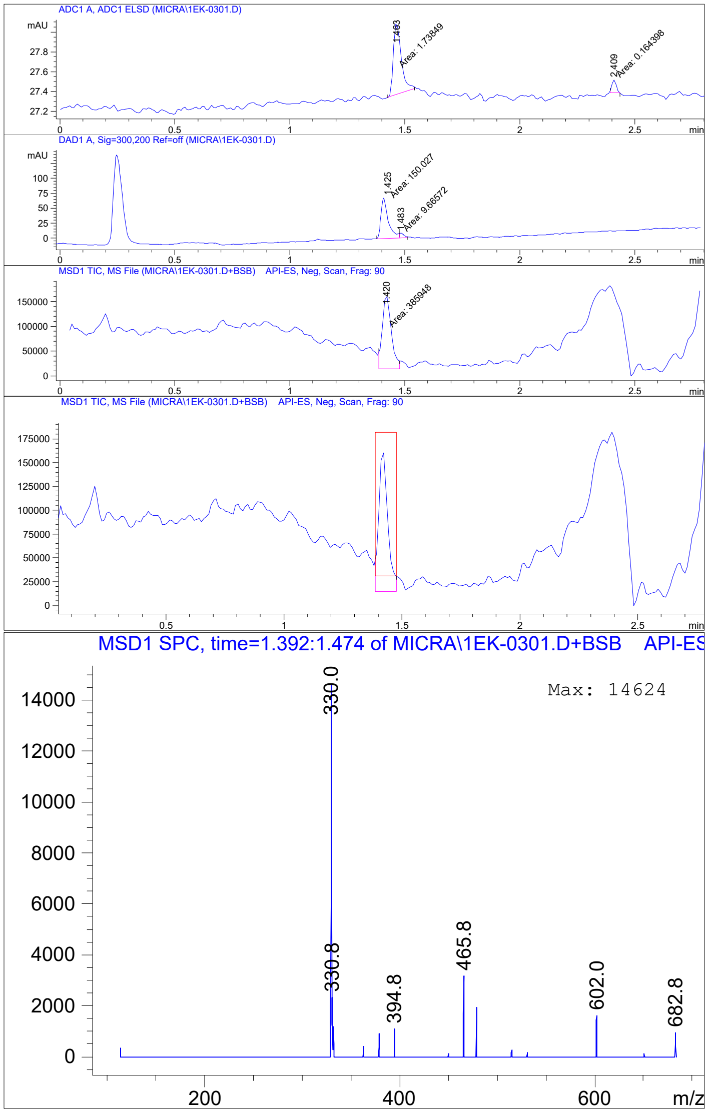

**In vitro ADME Study**

#### 1. EQUILIBRIUM SOLUBILITY

##### 1.1. Experimental Procedure

The equilibrium solubility of one test article was measured in pH 7.4 aqueous buffer. The buffer was prepared by combining 50 mL of 0.2 M KH2PO4 with 150 mL of H2O, and then adjusting to pH 7.4 with 10 N NaOH. At least 1 mg of powder for each test article was combined with 1 mL of buffer to make a ≥ 1 mg/mL mixture. These samples were shaken on a Thermomixer® overnight at room temperature. The samples were then passed through a 0.45 μm PTFE syringe filter. The filtrate was sampled and diluted in duplicate 10-, 100-, 1000-, and 10000-fold into a mixture of 1:1 buffer:acetonitrile (ACN) prior to analysis. All samples were assayed by LC-MS/MS using electrospray ionization against standards prepared in a mixture of 1:1 assay buffer:ACN. Standard concentrations ranged from 1.0 μM to 0.3 nM. Analytical conditions are outlined in Appendix 1.

##### 1.2. Experimental Results

| Test Article | Solubility in Phosphate Buffer (μM) | | |
|---|---|---|---|
| | R1 | R2 | AVG |
| NBCoV1 | 4.35 | 5.08 | 4.72 |

#### 2. P-GP SUBSTRATE ASSESSMENT

##### 2.1. Experimental Procedure

Caco-2 cells (clone C2BBe1) were obtained from American Type Culture Collection (Manassas, VA). Cell monolayers were grown to confluence on collagen-coated, microporous membranes in 12-well assay plates. Details of the plates and their certification are shown below. The permeability assay buffer was Hanks' balanced salt solution (HBSS) containing 10 mM HEPES and 15 mM glucose at a pH of 7.4. The buffer in the receiver chamber also contained 1% bovine serum albumin. The dosing solution concentration was 5 µM of test article in the assay buffer

+/-- 1 µM valspodar. Cells were first pre-incubated for 30 minutes with HBSS containing +/- 1 µM valspodar. Cell monolayers were dosed on the apical side (A-to-B) or basolateral side (B-to-A) and incubated at 37°C with 5% $CO_2$ in a humidified incubator. Samples were taken from the donor and receiver chambers at 120 minutes. Each determination was performed in duplicate. The flux of lucifer yellow was also measured post-experimentally for each monolayer to ensure no damage was inflicted to the cell monolayers during the flux period. All samples were assayed by LC-MS/MS using electrospray ionization. Analytical conditions are outlined in Appendix 1. The apparent permeability (Papp) and percent recovery were calculated as follows:

$$Papp = (dC_r/dt) \times V_r/(A \times C_A) \qquad (1)$$

$$\text{Percent Recovery} = 100 \times ((V_r \times C_r^{final}) + (V_d \times C_d^{final}))/(V_d \times C_N) \qquad (2)$$

Where,

$dC_r/dt$ is the slope of the cumulative receiver concentration versus time in µM $s^{-1}$;

$V_r$ is the volume of the receiver compartment in $cm^3$;

$V_d$ is the volume of the donor compartment in

cm$^3$; A is t

he area of the insert (1.13 cm$^2$ for 12-well);
$C_A$ is the average of the nominal dosing concentration and the measured 120 minutedonor concentration in μM;
$C_N$ is the nominal concentration of the dosing solution in μM;

$C^{final}$ is the cumulative receiver concentration in μM at the end of the incubation period;
$C_d^{final}$ is the concentration of the donor in μM at the end of the incubation period.

Efflux ratio (ER) is defined as $P_{app}$ (B-to-A) / $P_{app}$ (A-to-B).

#### 2.2. Cell Batch Quality Control Results

| | | |
|---|---|---|
| Plates | 12-well | |
| Seed Date | 20Jan2021 | |
| Passage Number | 67 | |
| Age at QC (days) | 20 | |
| Age at Experiment (days) | 27 | **Acceptance Criteria** |
| Atenolol Papp, $10^{-6}$ cm/s | 0.193 | ≤ 0.5 |
| Propranolol Papp, $10^{-6}$ cm/s | 19.7 | 10-30 |
| Digoxin A-to-B Papp, $10^{-6}$ cm/s | 0.340 | N/A |
| Digoxin B-to-A Papp, $10^{-6}$ cm/s | 12.9 | N/A |
| Digoxin Efflux Ratio | 37.9 | ≥ 10 |

##### 2.3. Experimental results

| **Test Article** | **Direction** | **Recovery (%)** | **Papp ($10^{-6}$ cm/s)** | | | **Efflux Ratio** | **P-gp Substrate Classification** |
|---|---|---|---|---|---|---|---|
| | | | **R1** | **R2** | **AVG** | | |
| NBCoV1 | A-to-B | 46.2 | 18.1 | 15.6 | 16.9 | 1.21 | Negative |
| | B-to-A | 54.2 | 17.8 | 23.1 | 20.4 | | |
| NBCoV1 + 1 µM Valspodar | A-to-B | 34.4 | 20.4 | 20.6 | 20.5 | 1.15 | |
| | B-to-A | 56.6 | 20.9 | 26.4 | 23.6 | | |

P-gp Substrate Classification:

CER ≥ 1.0 without valspodar, and reduced by ≥ 50% with valspodar: **Positive**

CER ≥ 1.0 without valspodar, and reduced by < 50% with valspodar: **Negative**

CER < 1.0 without and with valspodar: **Negative**

CER = Corrected Efflux Ratio = ER – 1

### 3. PLASMA PROTEIN BINDING

#### 3.1. Experimental Procedure

Studies were carried out in mixed-gender human plasma (Lot# AS1650-36), obtained from BioIVT and collected on K2EDTA. A Pierce Rapid Equilibrium Dialysis (RED) device was used for all experiments. Stock solutions of the test article and control compound were first prepared in DMSO. Aliquots of the DMSO solutions were dosed into 1.0 mL of plasma at a dosing concentration of 5 μM for the test article and 10 μM for the co-dosed control compound, warfarin. Plasma (300 μL), containing test article and control compound, was loaded into two wells of the 96-well dialysis plate. Blank phosphate-buffered saline (PBS) (500 μL) was added to each corresponding receiver chamber. The device was then placed into an enclosed heated rocker that was pre-warmed to 37°C, and allowed to incubate for four hours. After 4 hours of incubation, both sides were sampled.

Aliquots (50 μL for donor, 200 μL for receiver) were removed from the chambers and placed into a 96-well plate. Plasma (50 μL) was added to the wells containing the receiver samples, and

200 μL of PBS was added to the wells containing the donor samples. Two volumes of acetonitrile (ACN) were added to each well, and the plate was mixed and then centrifuged at 3,000 rpm for 10 minutes. Aliquots of the supernatant were removed, diluted 1:1 into water, and analyzed by LC-MS/MS.

Protein binding values were calculated as follows:

% Bound = [(PARR in Donor – PARR in Receiver) / (PARR in Donor)] × 100 PARR = peak area response ratio to internal standard, including applicable dilution factors.

#### 3.2. Experimental Results

| Test Article | % Bound | | | | | |
|---|---|---|---|---|---|---|
| | Test Article | | | Warfarin | | |
| | R1 | R2 | AVG | R1 | R2 | AVG |
| NBCoV1 | > 99.5 | > 99.5 | > 99.5 | 98.4 | 98.2 | 98.3 |

Warfarin binding acceptance criteria: ≥ 98.0 % bound

#### 4. STABILITY IN LIVER MICROSOMES

##### 4.1. Experimental Procedure

Mixed-gender human liver microsomes (Lot# 1010420) were purchased from XenoTech. The reaction mixture, minus NADPH, was prepared as described below. The test article was added into the reaction mixture at a final concentration of 1 µM. The control compound, testosterone, was run simultaneously with the test article in a separate reaction. The reaction mixture (without cofactor) was equilibrated in a shaking water bath at 37°C for 5 minutes. The reaction was initiated by the addition of the cofactor, and the mixture was incubated in a shaking water bath at 37°C. Aliquots (100 µL) were withdrawn at 0, 15, 30, 60, 90, and 120 minutes. Test article and testosterone samples were immediately combined with 400 µL of ice-cold 50/50 acetonitrile (ACN)/H2O containing 0.1% formic acid and internal standard to terminate the reaction. The samples were then mixed and centrifuged to precipitate proteins. All samples were assayed by LC-MS/MS using electrospray ionization. Analytical conditions are outlined in Appendix 1. The peak area response ratio (PARR) to internal standard was compared to the PARR at time 0 to determine the percent remaining at each time point. Half-lives and clearance were calculated using GraphPad software, fitting to a single-phase exponential decay equation.

##### 4.2. Reaction Composition

Liver Microsomes 0.5 mg/mL

NADPH (cofactor) 1 mM Potassium Phosphate, pH 7.4 100 mM Magnesium Chloride5 mM

Test Article 1 µM

##### 4.3. Experimental Results

| Test Article | % Remaining of Initial (n=1) | | | | | | Half-life (min) | CLint (mL/min/ mg protein) |
|---|---|---|---|---|---|---|---|---|
| | 0 min | 15 min | 30 min | 60 min | 90 min | 120 min | | |
| NBCoV1 | 100 | 101 | 80.5 | 68.9 | 69.5 | 41.6 | 112 | 0.0124 |

| Control Compound | Species | Half-life (min) | CLint (ml/min/mg protein) | Acceptable Range (t1/2, min) |
|---|---|---|---|---|
| Testosterone | Human | 11.3 | 0.122 | ≤ 41 |

#### 5. CYP IC50

##### 5.1. Experimental Procedure

The test articles, at eight concentrations (0-10 µM), were incubated with pooled HLM (0.25 mg protein/mL) in phosphate buffer (100 mM, pH 7.4) containing $MgCl_2$ (5 mM), NADPH (1 mM), and an individual CYP probe substrate (at approximately Km). The reaction mixture minus NADPH was equilibrated in a shaking water bath at 37ºC for 5 minutes. The reaction was initiated by the addition of NADPH, followed by incubation at 37°C for 10-30 minutes depending on the individual CYP isoform. The reaction was terminated by the addition of two volumes of ice-cold acetonitrile. Negative (vehicle) controls were conducted without the test article. Positive controls were performed in parallel at a single concentration using known CYP inhibitors. After the removal of protein by centrifugation at 1640g (3000 rpm) for 10 minutes at 4ºC, the supernatants were transferred to a 96-well plate and diluted with water containing internal standard (stable isotope-labeled CYP probe metabolite). The formation of CYP probe metabolite was determined by LC-MS/MS.

#### 5.2. CYP Probe Substrates and Metabolites

| CYP | Probe Substrate | Metabolite | Incubation Time (min) | Positive Control Inhibitor |
|---|---|---|---|---|
| CYP1A2 | Phenacetin (63 μM) | Acetaminophen | 20 | α-Naphthoflavone (1 μM) |
| CYP2B6 | Bupropion (75 μM) | OH bupropion | 20 | Thio-TEPA (30 μM) |
| CYP2C8 | Amodiaquine (2 μM) | Desethylamodiaquine | 20 | Montelukast (5 μM) |
| CYP2C9 | Diclofenac (10 μM) | 4’-OH diclofenac | 20 | Sulfaphenazole (10 μM) |
| CYP2C19 | S-mephenytoin (40 μM) | 4’-OH mephenytoin | 30 | (+)-N-3-benzylnirvanol (5 μM) |
| CYP2D6 | Bufuralol (7 μM) | 1’-OH bufuralol | 20 | Quinidine (1 μM) |
| CYP3A | Midazolam (2.5 μM) | 1’-OH midazolam | 10 | Ketoconazole (1 μM) |
| | Testosterone (55 μM) | 6β-OH testosterone | 10 | Ketoconazole (1 μM) |

#### 5.3. Experimental Results

| CYP | % of Control Enzyme Activity (n=1)a | | | | | | | | IC50 (μM) |
|---|---|---|---|---|---|---|---|---|---|
| | 0 μM | 0.0137 μM | 0.0412 μM | 0.123 μM | 0.370 μM | 1.11 μM | 3.33 μM | 10 μM | |
| CYP1A2 | 100 | 99.2 | 96.7 | 93.8 | 92.2 | 89.8 | 69.5 | 41.3 | 7.40 |
| CYP2B6 | 100 | 97.5 | 98.1 | 90.4 | 94.2 | 76.9 | 47.0 | 15.2 | 3.19 |
| CYP2C8 | 100 | 102 | 89.0 | 98.3 | 87.8 | 76.9 | 25.3 | 3.87 | 2.08 |

| | | | | | | | | | |
|---|---|---|---|---|---|---|---|---|---|
| CYP2C9 | 100 | 99.4 | 98.2 | 97.2 | 95.1 | 85.1 | 59.9 | 29.8 | 5.01 |
| CYP2C19 | 100 | 103 | 110 | 101 | 102 | 92.0 | 70.8 | 40.7 | 7.31 |
| CYP2D6 | 100 | 95.8 | 98.7 | 98.6 | 101 | 97.0 | 84.3 | 59.2 | > 10 |
| CYP3A (Midazolam) | 100 | 96.7 | 97.0 | 94.5 | 95.1 | 95.9 | 88.2 | 61.6 | > 10 |
| CYP3A (Testosterone) | 100 | 96.6 | 97.0 | 94.8 | 96.3 | 94.6 | 82.2 | 59.9 | > 10 |

[a] Percent of control enzyme activity = 100 × (Enzyme activity in the presence of TA / Enzyme activity in the absence of TA). Enzyme activity was calculated from the peak area ratio of CYP probe metabolite to internal standard by LC-MS/MS.

| **Positive Control Inhibitor** | **CYP** | **IC50 (μM)** |
|---|---|---|
| α-Naphthoflavone | CYP1A2 | 0.0164 |
| Thio-TEPA | CYP2B6 | 4.22 |
| Montelukast | CYP2C8 | 0.750 |
| Sulfaphenazole | CYP2C9 | 0.595 |
| (+)-N-3-benzylnirvanol | CYP2C19 | 0.324 |
| Quinidine | CYP2D6 | 0.0641 |
| Ketoconazole | CYP3A (Midazolam) | 0.0450 |
| Ketoconazole | CYP3A (Testosterone) | 0.0270 |

#### 6. APPENDIX 1. ANALYTICAL METHOD

Liquid Chromatography

Column: Waters ACQUITY UPLC BEH Phenyl 30 × 2.1 mm, 1.7 □m

M.P. Buffer: 25 mM ammonium formate buffer, pH 3.5 Aqueous Reservoir (A): 90% water, 10% buffer

Organic Reservoir (B): 90% acetonitrile, 10% buffer Flow Rate: 0.7 mL/minute

Gradient Program:

| Time (min) | % A | % B |
|---|---|---|
| 0.00 | 90 | 10 |
| 0.50 | 1 | 99 |
| 0.75 | 1 | 99 |
| 0.80 | 90 | 10 |
| 1.00 | 90 | 10 |

Total Run Time: 1.00 minute

Autosampler: 10 L injection volume

Strong Needle Wash: water/methanol/2-propanol:1/1/1; with 0.2% formic acid Weak Needle Wash: 0.1% formic acid in water

Seal Wash: 10% methanol, 90% water Mass Spectrometer

Instrument: PE SCIEX API 4000

Interface: Turbo Ionspray

Mode: Multiple reaction monitoring

Method: 1.0 minute duration Settings:

| Test Article | +/- | Q1 | Q3 | DP | EP | CE | CXP | IS |
|---|---|---|---|---|---|---|---|---|
| NBCoV1 | + | 470.1 | 104.9 | 112 | 10 | 47 | 15 | 5500 |

**TEM:** 500˚C; **CAD:** 7; **CUR:** 30; **GS1:** 50; **GS2:** 50

#### Additional Methods

**Enzyme assay**. Six NBCoV compounds including four with the higher viral inhibitory activity and 2 with no activity were tested against the 3 enzymes ß-lactamase, trypsin, and malate dehydrogenase (MDH). Additionally, the activity of the NBCoV compounds was investigated against the two luciferase enzymes NanoLuc and FLuc, which were used in the single-cycle antiviral assays. For all the assays, the compounds were tested at the higher dose of 2000 nM and less than 0.1% of DMSO was present in each assay.

For the ß-lactamase inhibitory assay, we used the colorimetric Beta-Lactamase Inhibitor Screening Kit (Sigma-Aldrich Co. LLC) by following the manufacturer's instructions. Briefly, the ß-lactamase was incubated with the NBCoV compounds for 10 minutes at 25 °C before addition of the substrate and the absorbance was measured at 490 nm in Tecan M1000 microplate reader.

For the trypsin inhibitory assay, we used the Trypsin Activity Colorimetric Assay Kit (Sigma-Aldrich Co. LLC) by following the manufacturer's instructions. Trypsin was incubated with the NBCoV compounds for 10 minutes at 25 °C before addition of the substrate and the absorbance was measured at 405 nm.

For the MDH inhibitory assay, we used the colorimetric Malate Dehydrogenase Assay Kit (Sigma-Aldrich Co. LLC) by following the manufacturer's instructions. MDH was incubated with the NBCoV compounds for 10 minutes at 37 °C before addition of the substrate and the absorbance was measured at 450 nm.

For the NanoLuc inhibitory assay, 293T\17 cells were plated at $2X10^4$ cells /well in a 96-well tissue culture plate and incubated overnight at 37 °C. The following day, the cells were transfected with 0.1 µg/well of the pNL4-3ΔEnv-NanoLuc expression vector. Following 24 h incubation, the cells were washed with PBS and lysed with 50 µL of the cell culture lysis reagent. The lysates were incubated with the NBCoV compounds for 10 minutes at 25 °C. As control, lysates were untreated or treated with the Intracellular TE Nano-Glo® Substrate/Inhibitor (Promega). The lysates were then transferred to a black plate and mixed with the same volume of Nano-Glo® Luciferase reagent. The luciferase activity was immediately measured with a Tecan SPARK.

For the FLuc inhibitory assay, 293T\17 cells were plated at 2X$10^4$ cells /well in a 96-well tissue culture plate and incubated overnight at 37 °C. The following day, the cells were transfected with 0.1 µg/well of the pNL4-3.Luc.R-E- expression vector. Following 24 h incubation the cells were washed with PBS and lysed with 50 µL of the cell culture lysis reagent. The lysates were incubated with the NBCoV compounds for 10 minutes at 25 °C. As control, lysates were untreated or treated with 100 µM resveratrol (Sigma-Aldrich Co. LLC). The lysates were then transferred to a white plate and mixed with the same volume of luciferase assay reagent. The luciferase activity was immediately measured with a Tecan SPARK.

**Colloidal aggregation study**

The cytotoxicity of NBCoV1 small molecule was evaluated in 293T/ACE2 cells in the presence or absence of 0.025% of Tween-80. Briefly, 1x$10^4$/well 293T/ACE2 cells were plated in a 96-well cell culture plate and incubated overnight. The following day, aliquots of escalating concentrations of the NBCoV1 were added to the cells and incubated at 37 °C. Following 48 h incubation, the MTS reagent was added to the cells and incubated for 4 h at 37 ºC. The absorbance was recorded at 490 nm.

For the infectivity assay in the presence or absence of 0.025% of Tween-80, 293T/ACE2 cells were infected with same amounts of pseudovirus pNL4-3ΔEnv-NanoLuc/SARS-CoV-2. After 48 h incubation, the cells were washed with PBS and lysed with 50 µL of lysis buffer. Twenty-five µL of the cell lysates were transferred to a white plate and mixed with the same volume of Nano-Glo® Luciferase reagent. The luciferase activity was measured immediately with the Tecan SPARK.

The antiviral activity of small molecules NBCoV1 in 293T/ACE2 cells infected with pseudoviruses SARS-CoV-2 was evaluated following the compound's centrifugation. Briefly, a concentrated aliquot of NBCoV1 small molecule in PBS solution was spun at 13000 g for 20 min as previously described (1), and the resulting supernatant was used for the neutralization experiment. For direct comparison, the assay was ran in parallel by using NBCov1 which was not spun. Briefly, 293T/ACE2 cells were plated at $1x10^4$/well in 96-well plates and incubated overnight. On the following day, aliquots of the pseudovirus at about 1500 $TCID_{50}$/well at an MOI of 0.1 were pretreated with graded concentrations of the NBCoV1-centrifuged or not (NBCoV1-control) for 30 min and added to the cells. 293T/ACE2 cells cultured with medium with or without the SARS-CoV-2 pseudovirus, were included as positive and negative controls, respectively. After 48 h incubation, the cells were washed with PBS and lysed with 50 *µ*L of lysis buffer. Twenty-five µL of the cell lysates were processed as above to measure the luciferase activity and calculate the percent inhibition by the NBCoV small molecules and the $IC_{50}$ values.

**Table-S1.** Antiviral activity of NBCoV small molecules against pseudovirus NL4-3ΔEnv-NanoLuc/SARS-CoV-2 ($IC_{50}$) in single-cycle assay in 293T/ACE2 cells (cells were pre-treated with the NBCoV compounds for 30 min before infection) and antiviral activity of NBCoV small molecules against the replication-competent authentic virus SARS-CoV-2 (US_WA-1/2020) virus ($IC_{100}$) in Vero E6 cells (cells were pre-treated for 2 h before infection).

| Compound | Pre-treated 293T/ACE2 cells/ SARS-CoV-2 $IC_{50}$ (nM) | Pre-treated Vero cells/ SARS-CoV-2 $IC_{100}$ (μM) |
|---|---|---|
| NBCoV1 | >2000 | >10 |
| NBCoV2 | >2000 | >10 |
| NBCoV3 | >2000 | >10 |
| NBCoV4 | >2000 | >10 |
| NBCoV5 | >2000 | >10 |
| NBCoV6 | >2000 | >10 |
| NBCoV7 | >2000 | >10 |
| NBCoV8 | >2000 | >10 |
| NBCoV9 | >2000 | >10 |

**Table-S2**. Evaluation of NBCoV small molecules against ß-lactamase, Trypsin, and MDH

| Compound | ß-lactamase $IC_{50}$ (nM) | Trypsin $IC_{50}$ (nM) | MDH $IC_{50}$ (nM) |
|---|---|---|---|
| NBCoV1 | >2000 | >2000 | >2000 |
| NBCoV2 | >2000 | >2000 | >2000 |
| NBCoV3 | >2000 | >2000 | >2000 |
| NBCoV4 | >2000 | >2000 | >2000 |
| NBCoV5 | >2000 | >2000 | >2000 |
| NBCoV34 | >2000 | >2000 | >2000 |

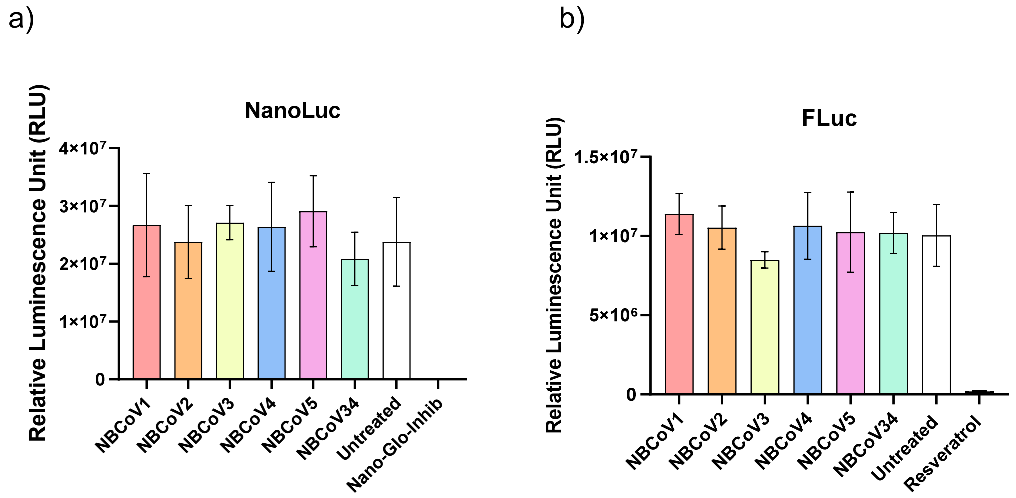

**Figure S1. Evaluation of NBCoV small molecules against NanoLuc and FLuc enzymes.** Lysates of 293T\17 cells expressing a) NanoLuc or b) FLuc were incubated with the NBCoV compounds for 10 minutes at 25 °C. As control, lysates were untreated or treated with the Intracellular TE Nano-Glo® Substrate/Inhibitor a) or with 100 µM of Resveratrol b). The lysates were then, mixed with the same volume of the respective substrate. Columns represent the means ± standard deviations (n = 2-3)

**The molecule with SMILES**
**OC(=O)C1=CC(=CC=C1Cl)C1=CC=C(O1)\C=C1/SC(=S)N(CCC2=CC=CC=C2)C1=O**
**OC(=O)C1=CC(=CC=C1)C1=CC=C(O1)\C=C1/SC(=S)N(CCC2=CC=CC=C2)C1=O**
**OC(=O)C1=CC(=CC=C1F)C1=CC=C(O1)\C=C1/SC(=S)N(CCC2=CC=CC=C2)C1=O**
**CC1=CC=C(C=C1C(O)=O)C1=CC=C(O1)\C=C1/SC(=S)N(CCC2=CC=CC=C2)C1=O**
**CC1=C(C=CC=C1C1=CC=C(O1)\C=C1/SC(=S)N(CCC2=CC=CC=C2)C1=O)C(O)=O**
**CCC1=CC=C(C=C1C(O)=O)C1=CC=C(O1)\C=C1/SC(=S)N(CCC2=CC=CC=C2)C1=O**
**OC(=O)C1=C(F)C(=CC=C1)C1=CC=C(O1)\C=C1/SC(=S)N(CCC2=CC=CC=C2)C1=O**
**OC(=O)C1=CC(=CC=C1Br)C1=CC=C(O1)\C=C1/SC(=S)N(CCC2=CC=CC=C2)C1=O**
**CC1=C(C=NC=C1C1=CC=C(O1)\C=C1/SC(=S)N(CCC2=CC=CC=C2)C1=O)C(O)=O**
**ClC1=CC=C(C=C1)C1=CC=C(O1)\C=C1/SC(=S)N(CCC2=CC=CC=C2)C1=O**
**S=C(S/1)NC(C1=C\C(O2)=CC=C2C3=CC=C(Cl)C(C(O)=O)=C3)=O**
**S=C(S/1)NC(C1=C\C(O2)=CC=C2C3=CC=C(C(O)=O)C(Cl)=C3)=O**
**S=C(S/1)N(CC#C)C(C1=C\C(O2)=CC=C2C3=CC=C(Cl)C(C(O)=O)=C3)=O**
**S=C(S/1)NC(C1=C\C(O2)=CC=C2C3=CC=C(C(O)=O)C=C3)=O**

was not similar to any known aggregator in our database.

However, since this molecule has a fairly high calculated LogP of **5.1** which is in the range reported for many other aggregators, we strongly urge you to perform appropriate controls to test for possible aggregation. Click here for guidance on how to perform these tests.

Created by Gurgen Tumanian, Allison Doak, Teague Sterling, John Irwin and Brian Shoichet with assistance from Pascal Wassam. We thank NIGMS for financial support. top

**Figure S2.** The data from Aggregator Advisor run could not find any known aggregator with similar structure.

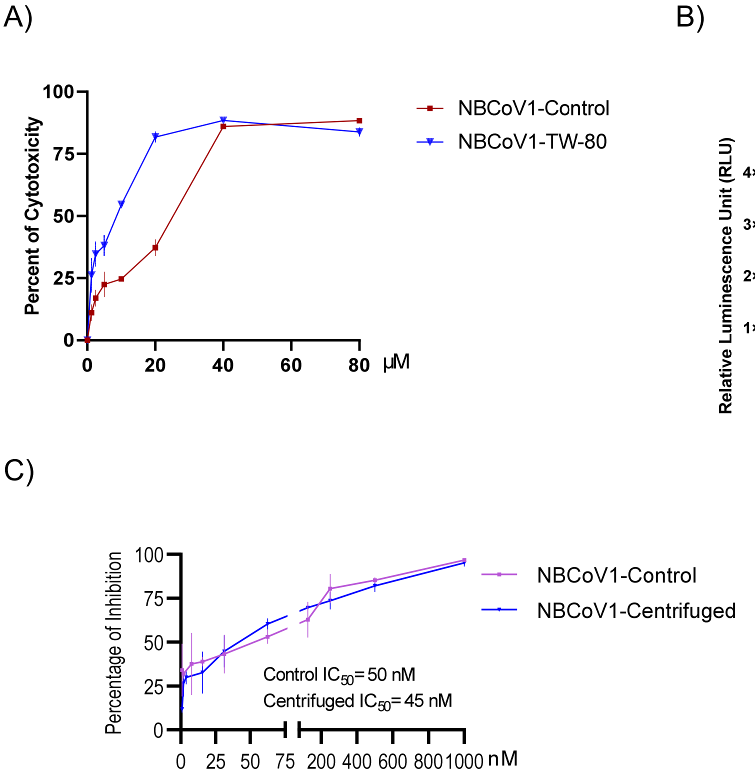

**Figure S3. Colloidal-aggregation studies with NBCoV1**. A) Cytotoxicity assay of NBCoV1 performed in 293T/ACE2 cells in the presence or absence of 0.025% of Tween-80 (the values represent the means ± standard deviations (n = 3)). B) Infection of 293T/ACE2 cells with same amounts of pseudovirus SARS-CoV-2 in the presence or absence of 0.025% of Tween-80 (control). Columns represent the means ± standard deviations (n = 3). C) Neutralization assay with NBCoV1-centrifuged and not-centrifuged (NBCoV1-control) in 293T/ACE2 cells infected with pseudovirus SARS-CoV-2 (the values represent the means ± standard deviations (n = 3).
